# Persistent but variable effect of experimental laboratory burns on microbial community resistance, resilience, and function across contrasting boreal forest soils

**DOI:** 10.64898/2026.09.01.748615

**Authors:** Dana B. Johnson, Kara M. Yedinak, Thea Whitman

**Affiliations:** University of Wisconsin-Madison, Madison, Wisconsin, U.S; Forest Products Laboratory, USDA Forest Service, Madison, Wisconsin, U.S; The University of British Columbia, Vancouver, British Columbia, Canada

**Keywords:** boreal forest soil, soil microbial community, wildfire, resistance, resilience, carbon use efficiency, gene copy number

## Abstract

Boreal forests stretch across vast swaths of the northern hemisphere, are shaped by wildfire, and play an important role in the global carbon cycle. Microorganisms play a critical role in soil nutrient cycling in these ecosystems, yet there are many open questions about the impacts of wildfire on microbially mediated soil biogeochemical cycles. In this study, we used laboratory burns and soil incubations of intact soil cores collected from two distinct soil types – Histosols and Gleysols – from boreal forest within Wood Buffalo National Park, Alberta, Canada, to assess burn effects on soil bacterial and fungal community composition and function. We compared resistance and resilience to burning for microbial communities vs. resistance and resilience to burning for soil pH and soil respiration to assess the relationships between burn-induced shifts in microbial community composition, the soil environment, and microbial activity. To link shifts in microbial community composition to potential community function, we measured glucose-specific carbon use efficiency (CUE) and assessed its relationship with weighted mean predicted 16S rRNA gene copy numbers for bacterial communities and FUNGuild-estimated relative abundance of putative symbiotrophic and saprotrophic fungi in burned and unburned soils. Microbial community resistance and resilience to burning varied across soil type with higher resistance of both bacterial and fungal communities from Histosols compared to the O horizons of Gleysols. This may be explained by a larger impact of burning on microbes in the thinner Gleysol O horizons. The relatively low resilience of bacterial and fungal communities to burning as well as the failure of resilience to increase with time since burning supports previous reports of post-burn microbial community recovery occurring over years rather than months. Burning caused a decrease in CUE with larger decreases following longer, hotter burns, which correlated with an increase in weighted mean predicted 16S rRNA gene copy number, raising the possibility that copy number could serve as a proxy for post-fire CUE in boreal forest soils, though more research is needed to constrain the effects of environmental conditions, substrates, and time since fire on this relationship. These findings suggest several ways in which burn-induced shifts in microbial community composition reflect altered microbial community function in meaningful ways for soil carbon cycling.

**Highlights:**

- Soil microbial community resistance and resilience to burning varies across boreal forest soil types
- No evidence of microbial community composition recovery within 70 days of fire
- Burning decreases glucose-specific CUE with larger decreases following longer, hotter burns
- Burning causes a rapid and persistent increase in weighted mean predicted 16S rRNA gene copy number

## 1. Introduction

Boreal forests are the world’s largest terrestrial biome and contain a globally important reservoir of carbon (C) both above- and belowground (Scharlemann et al., 2014; Bradshaw and Warkentin, 2015). These forests are well adapted to fire, having burned on 50 to 300 year fire return intervals for thousands of years (Kasischke et al., 1995; Stocks et al., 1996; Ter-Mikaelian et al., 2009; de Groot et al., 2013a, 2013b; Kelly et al., 2013; Prince et al., 2018). However, changes in regional patterns of temperature and precipitation appear to be driving a shift in fire regimes towards higher severity fires, shorter fire return intervals, and/or larger wildfires (Kasischke and Turetsky, 2006; Kasischke et al., 2010; de Groot et al., 2013b; Kelly et al., 2013; E. Whitman et al., 2022). Understanding how shifting climate and wildfires will impact boreal forest C cycling requires knowledge of controls on C stocks and decomposition rates. Soil microbes are important mediators of the global C cycle, and there is growing interest in soil microbial response to wildfires, which have burned 3% of the Earth’s terrestrial surface annually over the past 10 years (Global Wildfire Information System, 2026). A growing number of studies document shifts in soil microbial community composition following fire (e.g., Holden et al. 2016; Pérez-Valera et al. 2019; T. Whitman et al. 2019; Dove et al. 2022; Johnson et al. 2023, 2024); however, little is known about how soil microbial community composition and function will respond to shifts in boreal forest wildfire regimes towards hotter, larger fires. This is an important knowledge gap because understanding the response of soil microbial communities to current and future fire regimes is a critical component for predicting the fate of boreal forest soil C stocks in the coming decades. Current genome-based analyses of microbial communities are powerful, but only allow us to infer predicted functional potential (Cobo-Díaz et al., 2015; Dove et al., 2022; Nelson et al., 2022); thus, direct assessments of burn-induced shifts in soil microbial function are needed to better understand whether and how microbially mediated processes change soil organic C cycling post-fire. Additionally, our current understanding of microbial response to fire is temporally limited. With the exception of work taking advantage of wildfires burning pre-existing study sites (Glassman et al., 2016), and the work of a few intrepid researchers (Borgogni et al., 2019), most studies documenting the effects of burning and burn severity on post-fire microbial communities in forest soils begin months to years post-fire (Treseder et al., 2004; Waldrop and Harden, 2008; Sun et al., 2015; Holden et al., 2016; Dove et al., 2022; Nelson et al., 2022; T. Whitman et al., 2022; Lou et al., 2023) and are not able to capture shorter term soil microbial community dynamics. Predicting microbial response to shifting wildfire regimes may be aided by a better understanding of the short-term (days to months) dynamics of post-burn soil microbial communities.

There are multiple mechanisms by which fire could affect soil microbes both immediately and in the weeks to months post-burn, including via high soil temperatures during the burn itself (DeBano, 1991; Neary et al., 1999), shifts in soil pH (Certini, 2005; Úbeda et al., 2005; Bodí et al., 2014; Masyagina et al., 2016; Dove et al., 2020; Luo, 2023; Johnson et al., 2024) and the loss of organic matter (Kasischke and Johnstone, 2005), all of which have documented effects on soil microbial communities (Lauber et al., 2009; Rousk et al., 2010; Pingree and Kobziar, 2019; Whitman et al., 2019; Dove et al., 2020). Below the soil surface, the immediate effect of fire is lessened due to the insulative properties of soil; accordingly, soil temperature rapidly decreases with increasing depth (Beadle, 1940). Thus, mass microbial mortality, organic matter combustion, and burn-induced alterations in the soil environment are expected to be greatest in the uppermost soil horizons. This is supported by previous research finding an immediate (48 hours) post-fire decrease in DNA and RNA concentrations (Johnson et al., 2023) and larger increases in soil pH in O horizon - but not underlying mineral - soil (Johnson et al., 2024).

Fire can also affect soil microbial communities via structural and chemical changes in C substrates. For example, microbes and plants that are killed but not combusted during a fire increase the soil necromass pool (Ribeiro-Kumara et al., 2022) and may become a nutrient source for surviving microbes. This necromass may create a niche for fast-growing taxa, leading to an enrichment of these fast-growers after fire. While it is difficult to quantitatively measure soil microbial community growth rates, 16S rRNA gene copy numbers offer a useful proxy for identifying fast-growing microbial taxa in burned communities (Yano et al., 2013; Roller et al., 2016). Increased 16S rRNA gene copy numbers have been documented in burned soils months to years post-fire (Nemergut et al., 2015; Whitman et al., 2019; Johnson et al., 2023), and have been linked to an increase in the relative abundance of experimentally identified fast-growing bacterial taxa following natural wildfires (Johnson et al., 2023). The proliferation of fast-growing taxa in burned soil may alter microbial community functioning if fast-growing taxa switch C sources or have lower carbon use efficiency (CUE). CUE is an estimate of the proportion of total substrate C taken up by microbes that is converted into microbial biomass C vs. respired as CO_2_ (Geyer et al., 2016). In a study of the effects of heating events on CUE, Yin et al. (2026) documented lower CUE in soils heated to 50°C compared to soils heated to 30°C or 40°C, which suggests that the effects of high temperature alone may alter CUE. Notably soil temperatures during wildfire can exceed hundreds of degrees Celsius (DeBano, 1991; Neary et al., 1999). Nelson et al. (2024) previously posited that fire-induced changes in CUE may have important implications for post-fire soil C stocks, but there is minimal published work on the relationship between burning and soil CUE. We were only able to find one study that measured the impact of wildfire on soil CUE (Auwal et al., 2023), which used C:N ratios and stoichiometry to predict CUE and found lower predicted microbial CUE in burned vs. unburned soils from forest sites across China. Decreasing CUE can be driven by a decrease in C allocated to microbial biomass and/or an increase in C respired as CO_2_ per unit substrate C metabolized. Notably, depressed soil respiration is frequently observed in post-burn burn soils (Bárcenas-Moreno et al., 2011; Holden et al., 2016; Kelly et al., 2021; Johnson et al., 2023, 2024). If the observed burn-induced decreases in soil respiration are driven by a reduction in C respired as CO_2_ per unit substrate C metabolized, then we would expect CUE to increase in post-burn soils. On the other hand, a consistent decrease in CUE in burned soils would indicate either decreased allocation of C to microbial biomass and/or increased C respiration per unit substrate C metabolized. Yin et al. 2026 documented decreased CUE in soils

Disentangling the impact of fire on soil microbial CUE is particularly important for improving model-based predictions of post-fire C emissions. C cycle models are an important tool in understanding the effects of ecosystem disturbances, such as wildfire, on soil C stocks, and CUE is a common parameter used in many C cycling models (Cotrufo et al., 2013; Sulman et al., 2018). Furthermore, there is strong interest in integrating disturbance-induced shifts in microbial community functioning, such as changes to maximum enzymatic rate (V_max_) and CUE, into existing C models to better study how disturbance events affect microbial functioning and soil C cycling. Currently, CUE is typically represented as a constant in C models, precluding any fire-induced shifts in CUE or timeline of recovery of CUE to pre-burn levels. While we acknowledge that a single lab study of substrate-specific CUE in burned soils is far from sufficient for informing CUE representation in existing soil models, exploring the potential for burning to alter soil CUE is an important step towards a more realistic representation of post-fire microbial dynamics in soil C models.

In this study, our objectives were to use metrics of resistance and resilience to assess the effects of burn conditions, soil type, and soil properties on microbial communities, and to explore ways in which burn-induced shifts in microbial community composition affect CUE and microbial respiration. Measuring the immediate effects of wildfires on soil communities is hindered by difficulties in accessing sampling sites in the days to weeks after a fire. To address this limitation, we collected unburned intact soil cores for both organic-rich (Histosols) and sandy soils (Gleysols) characteristic of boreal forests in northern Alberta, Canada, subjected them to burns in a laboratory setting to simulate boreal forest crown fire conditions, and set up laboratory incubations to measure the effects of fire on soil communities in the days to weeks post-burn.

We hypothesized that (1) the resistance and resilience of microbial community composition to burning would vary across soil types and burn treatment durations, with lower resistance and resilience in O horizon vs. mineral soils and in longer vs. shorter burn durations, due to higher burn temperatures, (2) burning would cause a shift towards faster growing, less efficient microbial communities, reflected in increased weighted mean predicted 16S rRNA gene copy numbers and decreased CUE, and (3) microbial community resilience to burning would increase and weighted mean predicted 16S rRNA gene copy number would decrease with time since burn due to recovery of microbial communities.

## 2. Methods

### 2.1. Overview

In brief, intact soil cores collected from Wood Buffalo National Park, Alberta, Canada were subjected to burn simulations in a mass loss calorimeter. After the burns, soils were incubated for 70 days. At four timepoints (2, 24, 49, and 70 days), replicate cores were destructively sampled, and horizon subsamples for each core were collected for soil property analysis (reported in Johnson et al. 2024) and DNA sequencing to assess the impact of burning on soil microbial communities over time. This study is a companion paper to Johnson et al. (2024), which analyzed heating profiles during the burns, tracked soil CO_2_ fluxes after burning over the course of the incubations, and measured soil pH and total C in soil cores for each timepoint.

### 2.2. Experimental design and soil collection

We collected intact soil cores from 12 sites within Wood Buffalo National Park, Alberta, Canada, that had not burned in the previous 30 years (Figures 1 & S1, Table S1). Six of the sites were primarily jack pine (*Pinus banksiana* Lamb.) underlain by sandy, acidic soils classified as Eutric Gleysols (Food and Agriculture Organization of the United Nations., 2003). The other six sites were primarily spruce (*Picea mariana* (Mill.) and/or *Picea glauca* (Moench) Voss) underlain by soils classified as Dystric Histosols (Food and Agriculture Organization of the United Nations., 2003). In June 2022, 18 soil cores 6 cm in diameter and 10 cm in height were collected from each of these 12 sites, and 12 cores were then randomly selected for the burn experiment and incubations described in this paper. For each site, these 12 cores were randomly assigned to two burn treatment groups and a control group, resulting in four cores for each treatment, with each core being destructively sampled at a different timepoint in the incubation.

**Figure 1.**
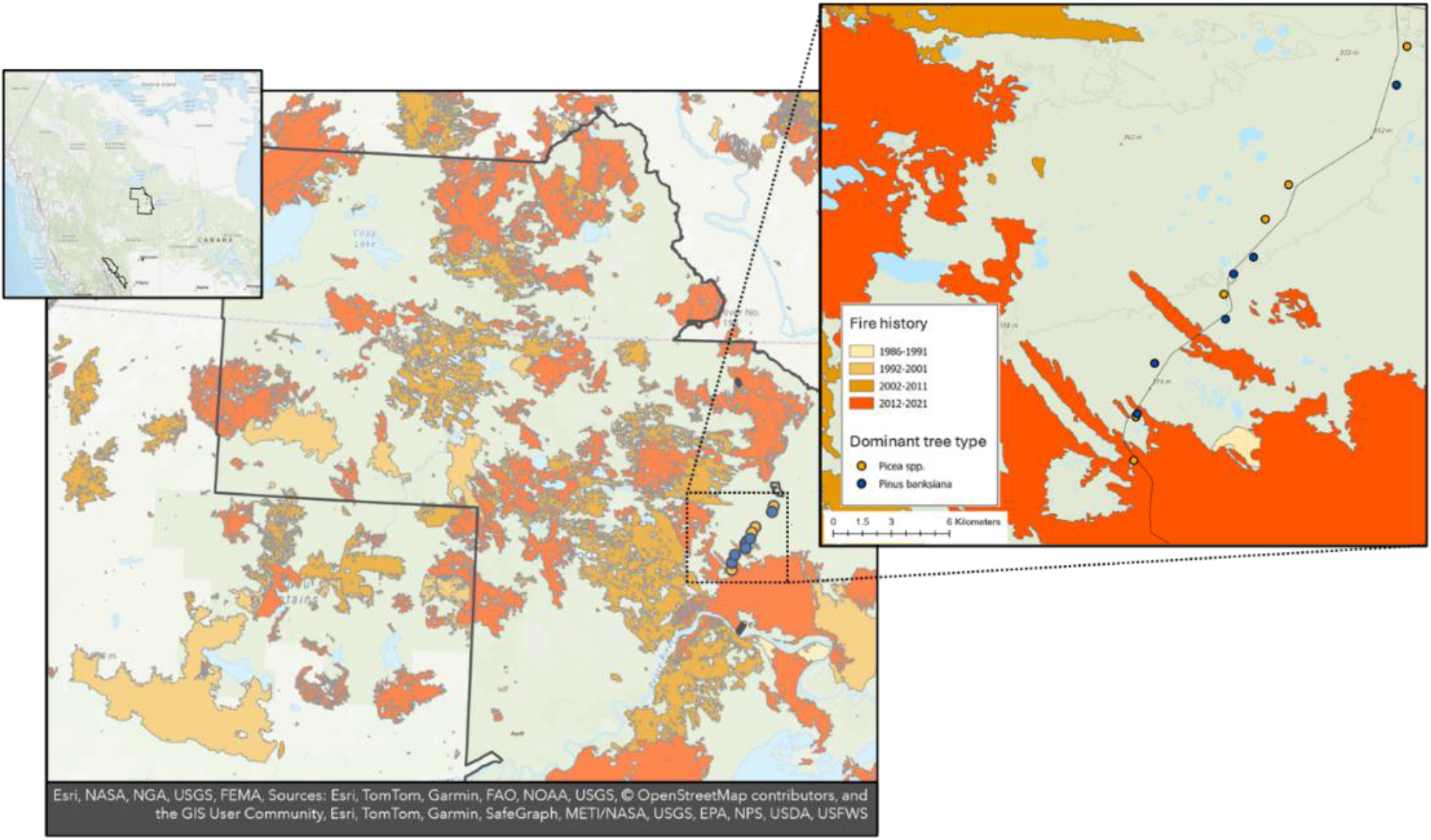
Soil cores were collected from 12 sites within Wood Buffalo National Park, Canada. Blue and yellow circles represent the location of sites dominated by Picea spp. and Pinus banksiana, respectively. The park boundary is outlined in black. Shades of orange and red mark areas burned since 1986 with darker colors denoting more recent fires.

Thus, for each timepoint, within a given soil/vegetation type, there are 6 samples, each representing an individual core from a separate site.

### 2.3. Laboratory burns, incubations, and sampling

To simulate burns, we exposed intact soil cores to a 60 kW m^-2^ heat flux in a Mass Loss Calorimeter (Fire Testing Technology Limited, West Sussex, UK) for either 30 s or 120 s, to mimic conditions at the soil surface during a mid- to high-range crown fire in the boreal forest (Frankman et al., 2012; Thompson et al., 2015) (see Johnson et al. (2024) for further details). After the burn, cores were allowed to cool to room temperature for 24 hours. The intact cores were incubated in capped 475 mL Mason jars for 2, 24, 49, or 70 days at a soil moisture of 65% of field capacity. Jars were opened daily to reduce the buildup of CO_2_ in the jar headspace.

After two days, the first set of cores – one control core and one core from each burn treatment from each site – was destructively sampled. For Gleysol cores, O horizon material and underlying mineral soil were separated and homogenized. For Histosol cores, all soil was organic, and thus, the entire core was homogenized. Subsamples of the Gleysol O horizon and mineral soil and Histosol organic soil were immediately collected and stored at -80 °C for later DNA extraction and sequencing. This was repeated 24, 49, and 70 days post-burn. At 24 days post-burn, additional soil subsamples were collected for measuring CUE. This timepoint was chosen to capture the short-term impact of burning on microbial community function while also allowing for microbial necromass created in the fire to be partially consumed.

### 2.4. DNA extraction, amplification, and sequencing

To assess change in community composition with time since burn, DNA was extracted from soil collected at 2, 24, 49, and 70 days post-burn along with empty-tube blanks, using DNeasy PowerLyzer PowerSoil kits (QIAGEN) following the manufacturer’s instructions.

DNA was amplified via triplicate PCR targeting the v4 region of the16S rRNA gene of bacteria and archaea with 515f and 806r primers (Walters et al., 2015) and targeting the ITS2 region of fungi with 5.8S-Fun and ITS4-Fun primers (Taylor et al., 2016) with barcodes and Illumina sequencing adapters added following Kozich et al. (2013) (all primers are listed in Tables S2 & S3).

PCR amplicon triplicates were pooled, purified, and normalized using a SequalPrep Normalization Plate Kit (ThermoFisher Scientific, Waltham, MA) following the manufacturer’s instructions. All samples and blanks were pooled, and library cleanup was performed using a Wizard SV Gel and PCR Clean-Up System (Promega, Madison, WI) following manufacturer’s instructions. The pooled library was sequenced using 2 x 300 paired-end Illumina MiSeq sequencing at the UW-Madison Biotechnology Center resulting in a total of 13M reads for 16S and 17M reads for ITS2.

### 2.5. Sequence processing and taxonomic assignments (including copy number calculation)

For 16S reads, we quality filtered, trimmed, dereplicated, learned errors, picked OTUs, and removed chimeras using DADA2 (Callahan et al., 2016) implemented in QIIME2 (Bolyen et al., 2019), resulting in 6.5M reads (mean 28,879 reads per sample). Samples with fewer than 1000 16S reads (5 samples total) were removed from the study. Taxonomy was assigned using the naïve Bayes classifier (Bokulich et al., 2018) in QIIME2 with the aligned 515f-806r region of the 99% OTUs from the SILVA database (SILVA 138 SSU) (Quast et al., 2013; Yilmaz et al., 2014; Glöckner et al., 2017). Archaea accounted for 0.3% of total reads, and we hereafter refer to 16S sequencing results as “bacterial communities” for brevity.

We calculated the mean predicted 16S rRNA gene copy number for each sample by first assigning a predicted mean 16S rRNA gene copy number to each genus in the study using the rrnDB RDP Classifier tool (v 2.14) (Stoddard et al., 2015). We calculated the mean copy number of all taxa with a genus-level assignment in the study and used this value as the predicted mean 16S rRNA gene copy number for taxa without a genus-level assignment. To calculate community-weighted mean predicted rRNA gene copy number, we calculated the relative abundance of each OTU after dividing the OTU counts by the predicted copy number. We then summed the product of the predicted copy number and the relative abundance of each OTU for all OTUs in each sample (Nemergut et al., 2015).

For ITS2 reads, we trimmed sequences using cutadapt (Martin, 2011). We quality filtered, trimmed, dereplicated, learned errors, picked OTUs, and removed chimeras using DADA2 (Callahan et al., 2016) implemented in QIIME2 (Bolyen et al., 2019) resulting in 12M reads (mean 52,283 reads per sample). Samples with fewer than 1000 ITS2 reads (9 samples total) were removed from the study. Taxonomy was assigned using the UNITE species hypothesis 99% threshold database version 10.0 (Abarenkov et al., 2024). We used the FUNGuild database (Nguyen et al., 2016) to assign ITS2 taxonomic assignments to functional groups. Following the recommendation of Nguyen et al. (2016), only guild assignments with a confidence level of “probable” or “highly probable” were included in our analyses. Of the 7345 OTUs in our study, 3858 were assigned to a functional guild FUNGuild, which accounted for an average of 73% (min. = 5%, max. = 99%) of ITS2 reads across samples.

### 2.6. Resistance and resilience

To assess the relationship between microbial community composition and the soil environment with increasing time since burning, we compared the resistance and resilience of soil microbial communities with soil pH resistance and resilience to burning. We also assessed the relationship between microbial community composition and function in the weeks to months post-burn by comparing microbial community resilience and soil respiration resilience to burning at 24, 49, and 70 days post-burn. We calculated metrics of resistance and resilience of soil pH and soil respiration, using data originally reported in Johnson et al. (2024) within each soil horizon and burn treatment duration following Orwin and Wardle (2004).

Resistance refers to the degree to which a given variable remains unchanged following burning. A unitless metric of resistance was calculated using unburned control soil cores and burned soil cores two days post-burn following Equation 1:

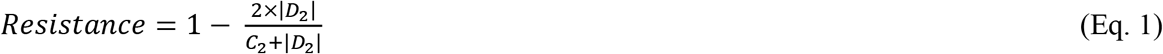

where D_2_ is the difference in a given variable for a burned core and the corresponding unburned control core from the same site two days after the burn, and C_2_ is the value of the variable for the unburned control core two days after the burn. Resistance can range from -1 to 1, with higher values representing less change with burning and lower values representing more change with burning.

Resilience refers to the degree of recovery of a given variable following burning and was calculated using unburned control soil cores and burned soils cores 24, 49, and 70 days post-burn following Equation 2:

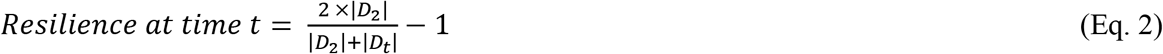

where *D_t_* is the difference in a given variable for a burned core and the corresponding unburned control core from the same site at time (*t*) since burn, and *D_2_* is as in Eq. 1 – the same metric two days after the burn.

We also calculated resistance and resilience of microbial community composition, informed by Sorensen and Shade (2020). The structure of the equations generally follows those above (Eqs. 1 and 2), but they draw on Bray-Curtis similarities, which are 1-Bray-Curtis dissimilarities (Bray and Curtis, 1957). Resistance was calculated from mean Bray-Curtis similarities of unburned soil cores and Bray-Curtis similarities of burned soils 2 days post-burn, following Equation 3:

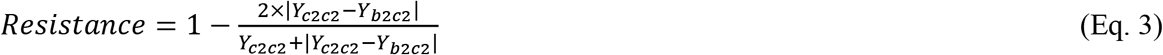

where *Y_c2c2_* is the mean Bray-Curtis similarity between a given control core and all other control cores 2 days post-burn (i.e., how similar on average is one site to the next?), and *Y_b2c2_* is the mean Bray-Curtis similarity between a given burned cores and corresponding control cores from all other sites 2 days post-burn (i.e., how similar on average is a burned core to an unburned core?). If burned community similarity to unburned communities is not less than site-to-site variability for unburned cores, then *Y_c2c2_* – *Y_b2c2_* will equal zero, and we will get a value of 1 – i.e., complete resistance. If the burned core communities are completely different from the control cores, *Y_b2c2_* will equal zero, so we get a value of 0 – i.e., no resistance at all.

We calculated resistance on day 2, as we expected this would best represent changes to community composition that were a direct result of the burn (i.e., due to combustion / cell death). Because DNA of dead organisms can persist in soils for some time, the truly optimal timepoint for this calculation would be some imaginary timepoint at which the DNA of all community members killed by fire had been degraded, but the composition of the living community members had not changed. Of course, this is impossible, so we limited our resistance calculation to the day two timepoint, with the recognition that it does likely include some taxa that were killed but for which their DNA has not yet degraded (thus potentially overestimating resistance).

To calculate resilience of microbial communities following burning, we used control communities two days into the incubation as the baseline value that the community could recover towards. The metric of resilience was thus calculated for communities at 24, 49, and 70 days post-burn using Bray-Curtis similarities between burned and unburned soils following Equation 4:

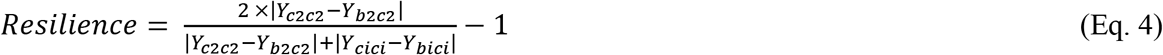

where *Y_c2c2_* and *Y_b2c2_* are as above in Eq. 3, and *Y_cici_* is the mean Bray-Curtis similarity between a control core at *i* days post-burn vs. all other control cores at *i* days post-burn, and *Y_bici_* is the mean Bray-Curtis similarity between a given burned core at *i* days post-burn vs. control cores from all other sites at *i* days post-burn. If burned core communities become more similar to control cores over time, *Y_cici_* – *Y_bici_* will approach zero and we will get a value of 1 – i.e., high resilience. If burned core communities diverge from control cores over time, *Y_cici_* – *Y_bici_* will increase yielding a negative value, and thus, lower resilience.

Metrics of resistance and resilience for soil pH, soil respiration (as total CO_2_ released per g C remaining for each incubation time period), and bacterial and fungal communities were calculated separately for each soil type, soil horizon, and burn treatment duration.

### 2.7. Carbon use efficiency

To assess the impact of burning on microbial community function, we measured CUE 24 days post-burn. After the 24-day incubation, the intact soil cores were destructively sampled. O and mineral horizons were separated, homogenized, and subsamples were collected for measurement of substrate-specific CUE using ^13^C-labelled glucose following the ^13^C-tracing method described in Geyer et al. (2019). Two subsamples were collected from each soil horizon and placed in 125 mL Mason jars. For Histosols, 1.5 g of soil was used, for Gleysol organic horizons, up to 2.0 g of soil was used, and for Gleysol mineral soils, 10 g of soil was used. The mass of soil used was limited by total intact horizon mass remaining after the burn. Thus, for six of the 18 soil horizons, less than 2.0 g of organic soil (0.7-1.3 g) was used. ^13^C-labelled glucose (5 at%) was then added to one of the jars and unlabeled glucose (1.1 at%) was added to the remaining jar at a rate of 1.0 mg glucose g^-1^ dry soil (0.4 mg C g^-1^ dry soil), (Table S4), and soil was mixed well.

These small rates of C addition were chosen with the goal of achieving detectable respiration and incorporation into microbial biomass while minimizing unrealistically large perturbances to the community due to new substrate additions. A parallel experiment testing burn effects on pine-specific CUE was run using labelled and unlabeled ground pine roots with inconclusive results, which are included in the supplementary data for completeness (Figure S2).

Jars were capped with lids fitted with rubber septa, sealed tightly, and incubated in the dark at 22 °C. A gas syringe was used to collect 12 mL headspace gas samples every 24 hours, and gas samples were injected into 12 mL exetainers and diluted 1:1 with CO_2_-free air for later analysis. Gas samples were also collected from additional empty jars to measure background CO_2_ levels. Following sampling, jars were opened, allowed to equilibrate with external atmosphere for five minutes, re-capped, and incubated for another 24 hours. A total of four headspace gas samples were collected from each jar over a 4-day period. Production and isotopic signature of CO_2_-C of headspace gas samples was measured by analyzing C concentration and isotopic composition on a cavity ring-down spectrometer (SAM 22001 autosampler; G2201-isotopic analyzer for CO_2_ and CH_4_, Picarro).

The degree of ^13^C-labelled glucose incorporated into microbial biomass was measured using a simultaneous chloroform fumigation extraction modified from Gregorich et al. (1990). Briefly, soil subsamples were placed in 20 mL of 0.05 M K_2_SO_4_ solution to which 0.5 mL ethanol-free chloroform was added (“fumigated”). 10× less concentrated K_2_SO_4_ was used than the original method so that C concentration in the salt solution would be sufficiently high so as to be detectable upon combustion (Potthoff et al., 2003; Makarov et al., 2015; Pang et al., 2021).

Duplicate samples with no chloroform were also used (“non-fumigated”). Solutions were shaken at 150 revolutions per minute for four hours after which solutions were filtered at 45 µm using a syringe filter, and the resulting dissolved organic carbon (DOC) was collected and stored. The DOC-K_2_SO_4_ solution was dried down at 80 °C for 24 hours, and the C concentration and isotopic composition of the resulting salts were quantified on a Thermo Delta V Isotope ratio mass spectrometer (IRMS) interfaced to a NC2500 elemental analyzer at the Cornell University Stable Isotope Laboratory.

Total microbial biomass C (MBC; µg C g^-1^ dry soil) was calculated using Equation 1 (see supplementary information for full list of equations and derivations):

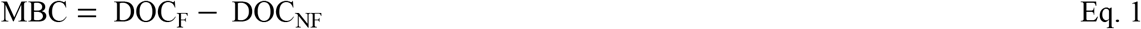

where DOC is the total C concentration of fumigated (F) and non-fumigated (NF) K_2_SO_4_ extracts on a per g dry soil basis. We did not correct MBC using an extraction efficiency coefficient (often termed k_EC_) because true extraction efficiencies vary widely across different soil properties (Dictor et al., 1998) and communities (Tate et al., 1988). Rather than multiplying all MBC estimates by the same value, which is almost certainly incorrect, we acknowledge that true MBC (and, hence, derived CUE) is presumably higher than our estimates by an unknown factor. Because we are primarily interested in comparing CUE from samples collected from the same soil type, which should have similar extraction efficiencies, this decision should not affect the direction of any observed trends.

Total microbial incorporation of ^13^C-labelled glucose C (MBC_LA_; µg C g^-1^ dry soil) was calculated as the product of MBC and the fraction of MBC that is glucose-derived (*f*_MBC,LA_) using Equations 2-5:

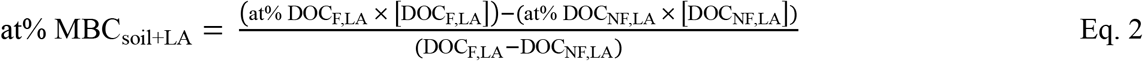

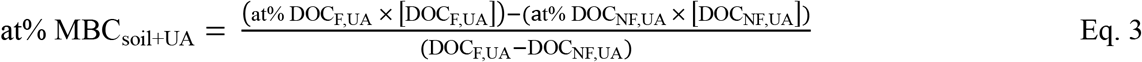

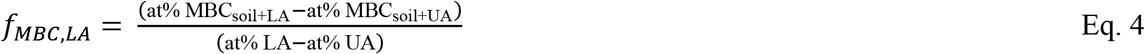

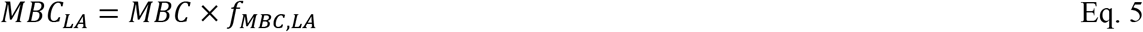

where at% MBC_soil_ and at% DOC represent the atom % of ^13^C in MBC and K_2_SO_4_ extracts, and DOC represents the concentration of C (µg C g^-1^ dry soil) of K_2_SO_4_ extracts from soils amended with labelled (LA) and unlabeled (UA) glucose, that were fumigated (F) or non-fumigated (NF). At% LA and at% UA are the atom % of ^13^C in the labelled and unlabeled glucose, respectively.

The total CO_2_ derived from the ^13^C-labelled glucose (CO_2,LA_; µg C g^-1^ dry soil) was calculated as the product of cumulative CO_2_-C respired (CO_2,total_; µg C g^-1^ dry soil) and the fraction of total CO_2_-C that is derived from the ^13^C-labelled glucose (*f*_CO2,LA_) using Equations 6-7:

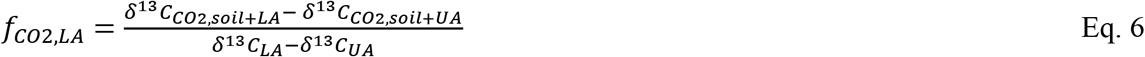

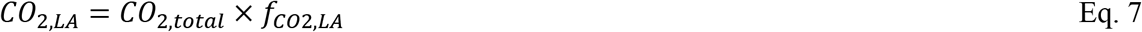

where δ^13^C_CO2,soil+LA_ and δ^13^C_CO2,soil+UA_ represent the δ^13^C values of CO_2_ from soils amended with the labelled (LA) or unlabeled (UA) glucose, respectively, and δ^13^C_LA_ and δ^13^C_UA_ represent the δ^13^C values of the labelled and unlabeled glucose, respectively.

Due to our relatively subtle addition of labelled glucose, glucose-derived MBC was not always detectable. Thus, before calculating CUE, we first filtered the data to include only datapoints with a positive value of substrate-derived MBC, which reduced the number of samples for glucose-amended soils from 54 to 53. CUE was then calculated using equation 8:

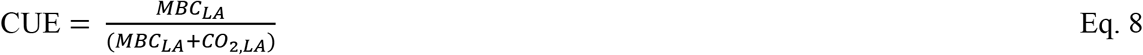

### 2.8. Bioinformatics and statistics

We performed all analyses using R statistical software version 4.4.0 (R Core Team, 2024), relying extensively on R packages phyloseq (McMurdie and Holmes, 2013), dplyr (Wickham et al., 2021), ggplot2 (Wickham, 2016), and vegan (Oksanen et al., 2020). We compared community composition across samples using Bray-Curtis dissimilarities (Bray and Curtis, 1957) on DNA-based relative abundances and tested for significant effects of burn treatment duration, controlling for soil type and horizon, soil pH, incubation length, and degree hours using a permutational multivariate ANOVA (PERMANOVA using the adonis function in vegan (Oksanen et al., 2020) and used the betadisper() function to test if variables differed in their dispersion. We used single-component models to compare the R^2^ for each factor.

We used ANOVA models and Tukey’s HSD to assess the impact of burning, burn treatment duration, and soil type on MBC, CO_2_ respired, and glucose-specific CUE (Oksanen et al., 2020). We performed nonlinear (exponential) regressions to evaluate the relationships between glucose- specific CUE and maximum soil temperature, soil pH, weighted mean predicted 16S rRNA gene copy number, and used linear regressions to evaluate the relationships between glucose-specific CUE and the relative abundance of putative symbiotrophic and saprotrophic fungi.

## 3. Results

### 3.1. Burn effects on microbial community composition

Burn treatments altered microbial community composition, with longer burn treatment durations resulting in larger shifts (Figures 2 & S3). All tested factors (burn treatment duration, burn degree-hours, soil type and horizon, pH, and incubation time) were significant predictors of community composition for both bacteria and fungi (PERMANOVAs, p < 0.01) (Tables S5-8). Soil type and horizon provided the most explanatory power for community composition for bacteria (R^2^ = 0.28; Figures S4-5) and fungi (R^2^ = 0.14; Figures S6-7), followed by pH (bacteria, R^2^ = 0.17; fungi, R^2^ = 0.06).

**Figure 2.**
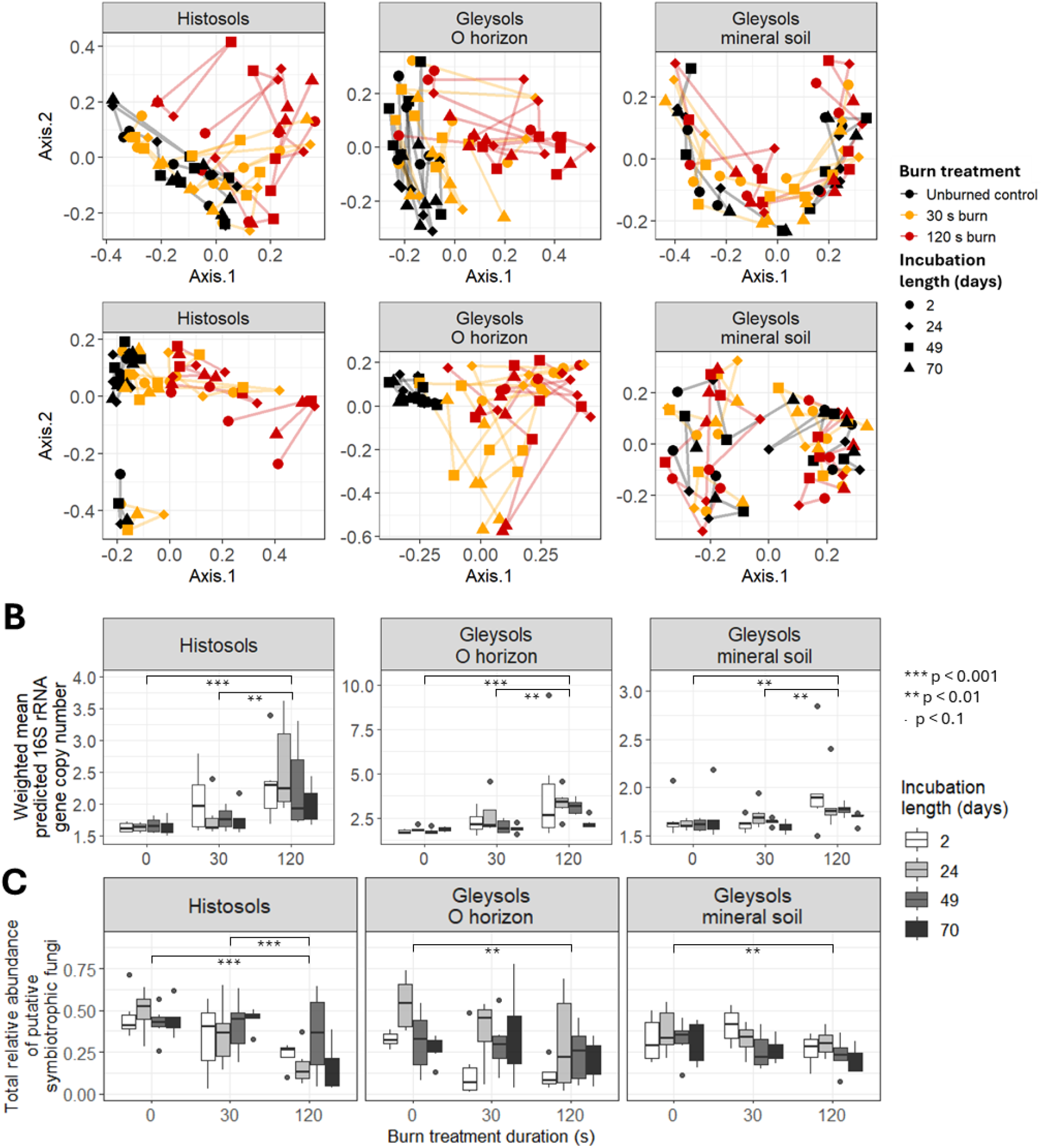
(A) Principal Coordinates Analysis (PCoA) of Bray-Curtis dissimilarities illustrating the effects of burn treatment duration on bacterial and archaeal communities (upper panels) and fungal communities (lower panels) across different soil horizons and incubation timepoints. Black, orange, and red represent soils exposed to 0, 30, and 120 s burn treatment durations, respectively, and circles, diamonds, squares, and triangles represent incubation duration of 2, 24, 49, and 70 days, respectively. (B) The effect of burning, burn treatment duration, and incubation length on weighted mean predicted16S rRNA gene copy number. (C) The effect of burning, burn treatment duration, and incubation length on the total relative abundance of putative symbiotrophic fungi. Statistically significant differences in burn treatment duration effects on weighted mean predicted16S rRNA gene copy number and the total relative abundance of putative symbiotrophic fungi are indicated on figure and based on ANOVAs and Tukey’s HSD. For boxplots, the central horizontal line indicates the median, the upper and lower bounds of the box indicate the inter-quartile range (IQR), the upper and lower whiskers reach the largest or smallest values with a maximum of 1.5xIQR, and data beyond the whiskers are indicated as individual points (n=6 except for the 120 s burn of Histosols at day 49 (n=5) and the 30 s burn of Gleysols at day 2 (n=5) and day 24 (n=5)).

Following burning, we observed a shift in bacterial communities towards higher weighted mean predicted 16S rRNA gene copy numbers in both Histosols and Gleysols (Figure 2; Tables S9- 10). The largest increase in weighted mean predicted 16S rRNA gene copy numbers was in soils exposed to the 120 s burn treatment duration. Elevated weighted mean predicted 16S rRNA gene copy numbers in burned vs. unburned soil persisted for the duration of the incubation in both Histosols and Gleysol O horizons.

While we did not assess analogous burn effects on predicted ITS2 copy number (due to fungal multicellularity and high intra- and interspecies variation in copy number (Lofgren et al., 2019; Bradshaw et al., 2023)), we used FUNGuild to screen for burn induced shifts in fungal trophic strategies. We observed a small decrease in the relative abundance of putative symbiotrophic fungi in Histosols and Gleysols exposed to the 120s burn treatment duration (Histosols, mean = 0.2 ± 0.2, p < 0.001; Gleysols, mean = 0.2 ± 0.2, p = 0.01) compared to unburned soil (Histosols, mean = 0.5 ± 0.1; Gleysols, mean = 0.4 ± 0.2) (Figure 2C). The fraction of the community classified as putative symbiotrophs was negatively correlated with weighted mean predicted 16S rRNA gene copy number in Histosols (y = -1.7x + 2.5, p < 0.001, R^2^_adj._ = 0.43) and O horizons of Gleysols (y = -2.4x + 2.5, *p* = 0.01, R^2^_adj._ = 0.07) (Figure S8). There was no significant difference in putative fungal saprotrophs or pathogens in burned vs. unburned soils; however, the fraction of the community classified as putative saptrotrophs was positively correlated with weighted mean predicted 16S rRNA gene copy number in O horizons of Gleysols (y = 2.3x + 2.0, *p* = 0.001, R^2^_adj._ = 0.13) and in Gleysol mineral soil (y = 0.81x + 1.6, *p* = 0.01, R^2^_adj._ = 0.08) (Figure S8).

### 3.2. Resistance

Microbial community resistance to burning ranged from 0.006 and 1.0, with bacterial communities showing higher resistance to burning (mean = 0.7 ± 0.2) than fungal communities (mean = 0.5 ± 0.3; ANOVA, *p* = 0.008) (Figure 3A and B). Microbial community resistance to burning varied across soil horizons but not between the two burn treatment durations. For example, the resistance of bacterial communities to burning was lower in Gleysol O horizons (mean = 0.5 ± 0.3) than in Gleysol mineral soil (mean = 0.8 ± 0.1; ANOVA, *p* = 0.01). Bacterial communities in Gleysols with thicker O horizons pre-burn showed marginally significant higher resistance to burning than communities in in soils with thinner O horizons (Figure S9; y = 0.21x + 0.02, R^2^_adj._ = 0.28, *p* = 0.06).

**Figure 3.**
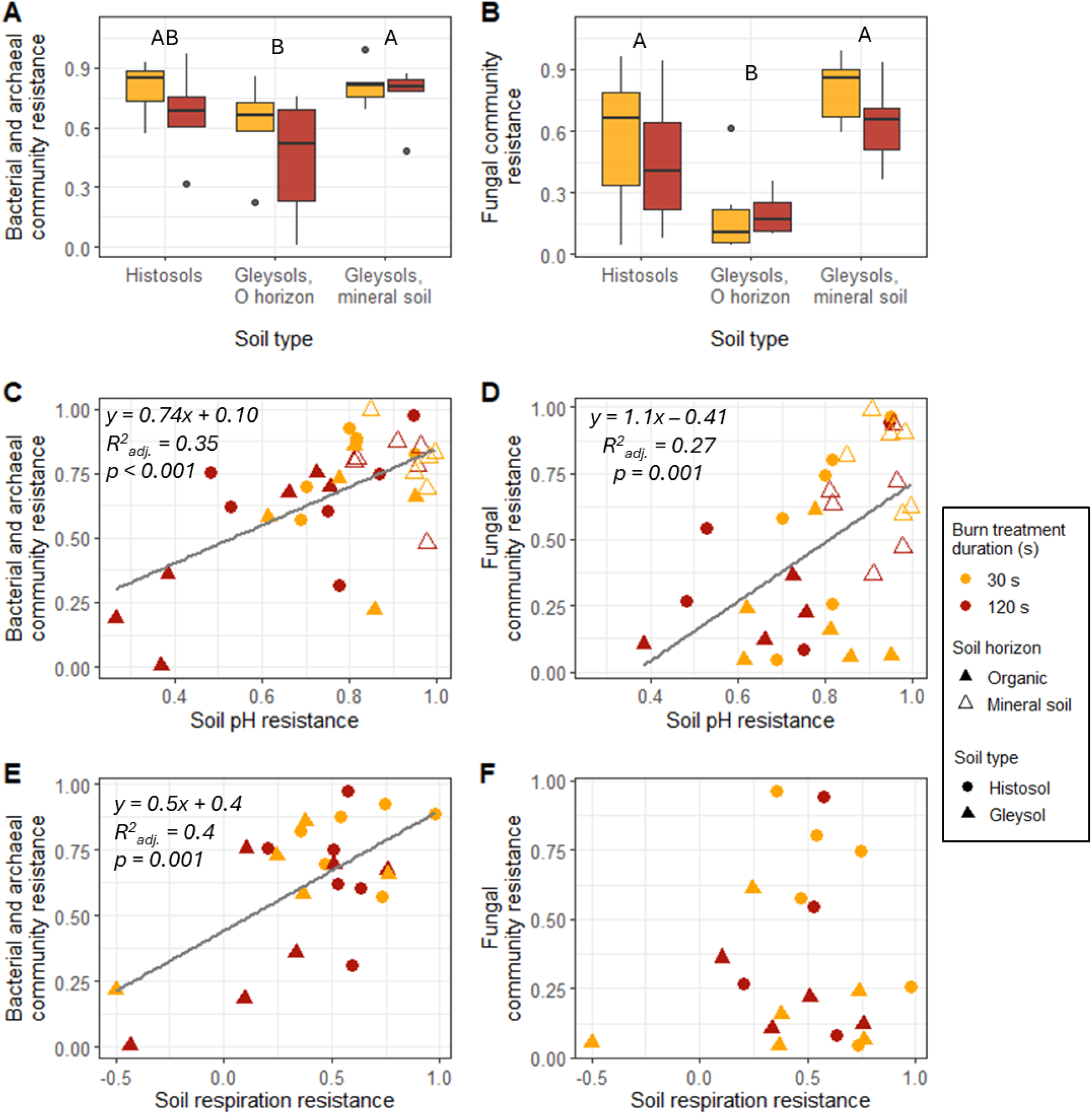
Resistance to burning of (A) bacterial and archaeal communities and (B) fungal communities across soil types and horizons. Different letters within a plot represent statistically significant differences between soil horizons based on ANOVA and Tukey’s HSD (p < 0.05). The central horizontal line indicates the median, the upper and lower bounds of the box indicate the inter-quartile range (IQR), the upper and lower whiskers reach the largest or smallest values within a maximum of 1.5 * IQR, and data beyond the whiskers are indicated as individual points. Resistance to burning of soil pH vs. (C) bacterial community resistance and (D) fungal community resistance. Resistance to burning of soil respiration vs. (E) bacterial community resistance and (F) fungal community resistance. Soil respiration resilience was calculated using cumulative CO_2_-C respired as a percent of C remaining post-burn. Orange and red represent soils exposed to the 30 s and 120 s burn treatment durations, respectively. Circles and triangles represent Histosols and Gleysols, respectively, with filled and unfilled symbols representing organic horizons and mineral soils, respectively.

The resistance of fungal communities to burning was lower in Gleysol O horizons (mean = 0.2 ± 0.2) than in mineral soil (mean = 0.7 ± 0.2; ANOVA, *p* < 0.001) or Histosols (mean = 0.5 ± 0.3; ANOVA, *p* = 0.02). Unlike with bacterial communities, we observed no significant effect of pre-burn O horizon thickness on fungal community resistance to burning in Gleysols (*p* = 0.13).

To assess the relationship between microbial community composition and burn effects on the soil environment, we compared the resistance to burning of microbial communities and soil pH and found that both bacterial and fungal community resistance was positively correlated with soil pH resistance to burning (Figure 3C & D). We also compared the resistance to burning of microbial communities and soil respiration (per g total C remaining post-burn) to assess the relationship between microbial community composition and function. The resistance of bacterial communities but not fungal communities was positively correlated with soil respiration resistance (Figure 3E & F).

### 3.3. Resilience

Soil microbial community resilience ranged widely, from -1.0 to 1.0, but mean resilience was low (mean = 0.04 ± 0.5) (Figure 4A & B). Resilience did not differ significantly between fungal vs. bacterial communities, and the effects of burn duration were small and varied across soil horizons. For example, bacterial community resilience was lower in Gleysol O horizons following the 120 s (mean = -0.09 ± 0.3) vs. 30 s burn treatment duration (mean = 0.3 ± 0.4, ANOVA, *p* = 0.008), but this effect was not evident in Histosols or mineral soil. Microbial community composition did not recover consistently over the course of the incubation across the different soil types – i.e., there was no significant relationship between resilience and time since burn for bacterial (*p* = 0.51) or fungal communities (*p* = 0.58).

**Figure 4.**
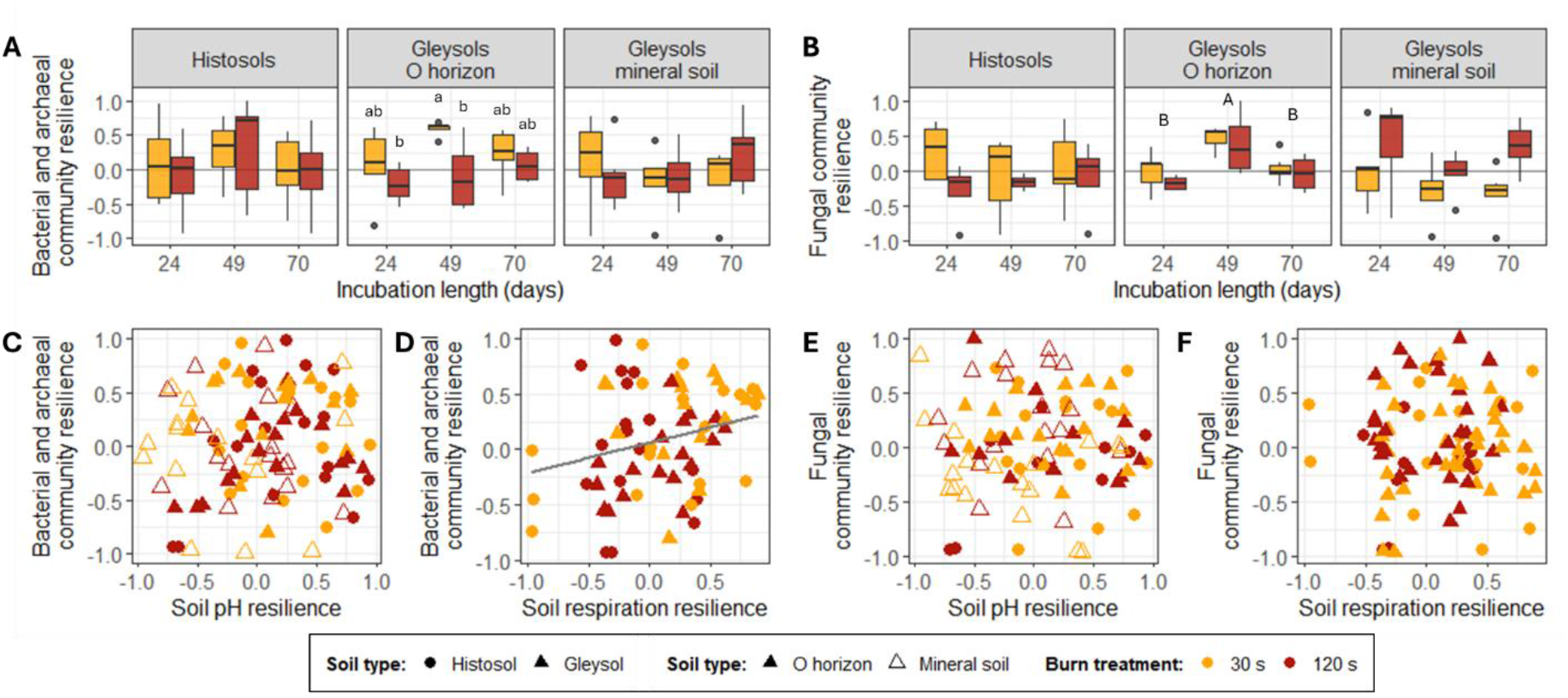
Resilience to burning of (A) bacterial and archaeal communities and (B) fungal communities across soil types and horizons. Different letters within a plot represent statistically significant differences between burn treatment duration (lowercase letters) or soil horizon (uppercase letters) based on ANOVA and Tukey’s HSD (p < 0.05). The central horizontal line indicates the median, the upper and lower bounds of the box indicate the inter-quartile range (IQR), the upper and lower whiskers reach the largest or smallest values within a maximum of 1.5 * IQR, and data beyond the whiskers are indicated as individual points. Resilience to burning of soil pH vs. (C) bacterial community resilience and (E) fungal community resilience. Resilience to burning of soil respiration vs. (D) bacterial community resilience (y = 0.3x + 0.06; R^2^_adj._ = 0.05; p = 0.04) and (F) fungal community resilience. Orange and red represent soils exposed to the 30 s and 120 s burn treatment durations, respectively. Circles and triangles represent Histosols and Gleysols, respectively, with filled and unfilled symbols representing organic horizons and mineral soils, respectively.

We were interested in the role of soil pH and burn-induced shifts in soil pH on microbial community resilience to burning, so we compared microbial community resilience to both pH resistance and resilience to burning. Microbial community resilience was not significantly correlated with pH resistance (bacteria, *p* = 0.72; fungi, *p* = 0.07) or pH resilience (bacteria, *p* = 0.49; fungi, *p* = 0.87) (Figure 4C & 4E).

Soils with a higher respiration resilience to burning, i.e., burned soils in which respiration rates (per g total C remaining post-burn) increased toward unburned soil respiration rates over time, also hosted bacterial communities with higher resilience (Figure 4D). There was no significant trend between respiration resilience and fungal community resilience (Figure 4F).

### 3.4. Glucose-specific CUE

To assess the effect of burning on microbial community function, we compared microbial community CUE in burned vs. unburned soils. CUE values ranged from 0.02 in burned soil to 0.5 in unburned soil (Figure 5C). There was no significant effect of soil type on the relationship between burning and CUE, and, thus, we report CUE by soil horizon, combining Gleysol O horizons and Histosols. Burning decreased CUE in organic horizons with lower CUE in soils exposed to the 120 s burn duration (mean = 0.08 ± 0.07, p < 0.001) than in soils exposed to the 30 s burn duration (mean = 0.16 ± 0.09, p < 0.001). Burning had no significant effect on CUE in mineral soils. The observed decrease in CUE in burned organic horizons was primarily driven by a decrease in the amount of amended glucose C incorporated into microbial biomass, with higher levels of glucose-derived MBC in unburned organic horizons (mean = 48 ± 14 µg g^-1^ dry soil) than in burned soils (30s burn treatment duration, mean = 22 ± 11 µg g^-1^ dry soil, p < 0.001; 120 s burn treatment duration, mean = 11 ± 8 µg g^-1^ dry soil, p < 0.001) (Figure 5B). In comparison, there was a small increase in glucose-derived CO_2_ from burned soils (120 s burn treatment duration: 129 ± 9 µg g^-1^ dry soil) compared to unburned soils (102 ± 24 µg g^-1^ dry soil, p = 0.008) (Figure 5A). The small mass of added glucose (Table S4) had little overall effect on MBC or respiration, as desired: the bulk of MBC and CO_2_ emitted over the four day incubation was derived from SOC and not from the added glucose in organic horizons (Figures S11-12). Total MBC and CO_2_ emitted from mineral soils was much lower than from organic soils, and a larger fraction of total MBC and CO_2_ was derived from added glucose (Figures S11-12).

**Figure 5.**
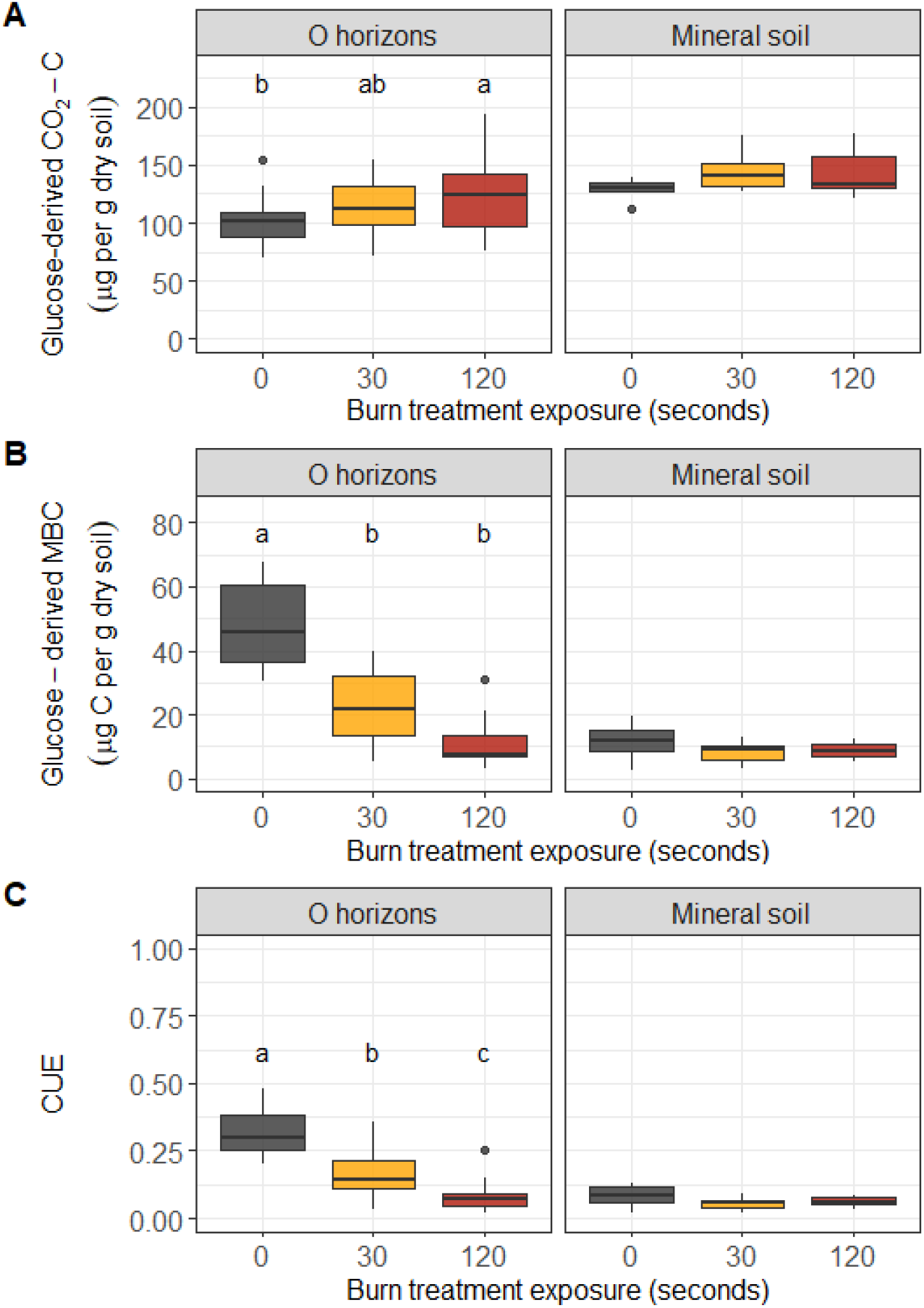
(A) Glucose-derived CO_2_-C, (B) glucose-derived microbial biomass C (MBC), and (C) glucose-specific CUE in organic (left panels) and mineral (right panels) soils. Different letters within a plot represent statistically significant differences burn treatment duration based on ANOVA and Tukey’s HSD (p < 0.05). The central horizontal line indicates the median, the upper and lower bounds of the box indicate the inter-quartile range (IQR), the upper and lower whiskers reach the largest or smallest values within a maximum of 1.5 * IQR, and data beyond the whiskers are indicated as individual points.

To assess the relationship between soil temperature during a burn and microbial community function, we compared the maximum soil temperature reached in each horizon vs. CUE. In organic horizons, CUE decreased exponentially with increasing maximum soil temperatures during the burn treatment (y = 0.6e^-0.04x^ + 0.07; p < 0.001, R^2^_adj._ = 0.62) (Figure 6A). There was weak evidence for a negative relationship between CUE and maximum soil temperature in the mineral horizons (y = 0.33e^-0.08x^ + 0.04; p = 0.06, R^2^_adj._ = 0.12), but the range of mineral soil temperatures was much more constrained (ranging from 25°C in unburned soils to about 55°C in burned soils) compared to temperatures in organic horizons (Figure 6A).

**Figure 6.**
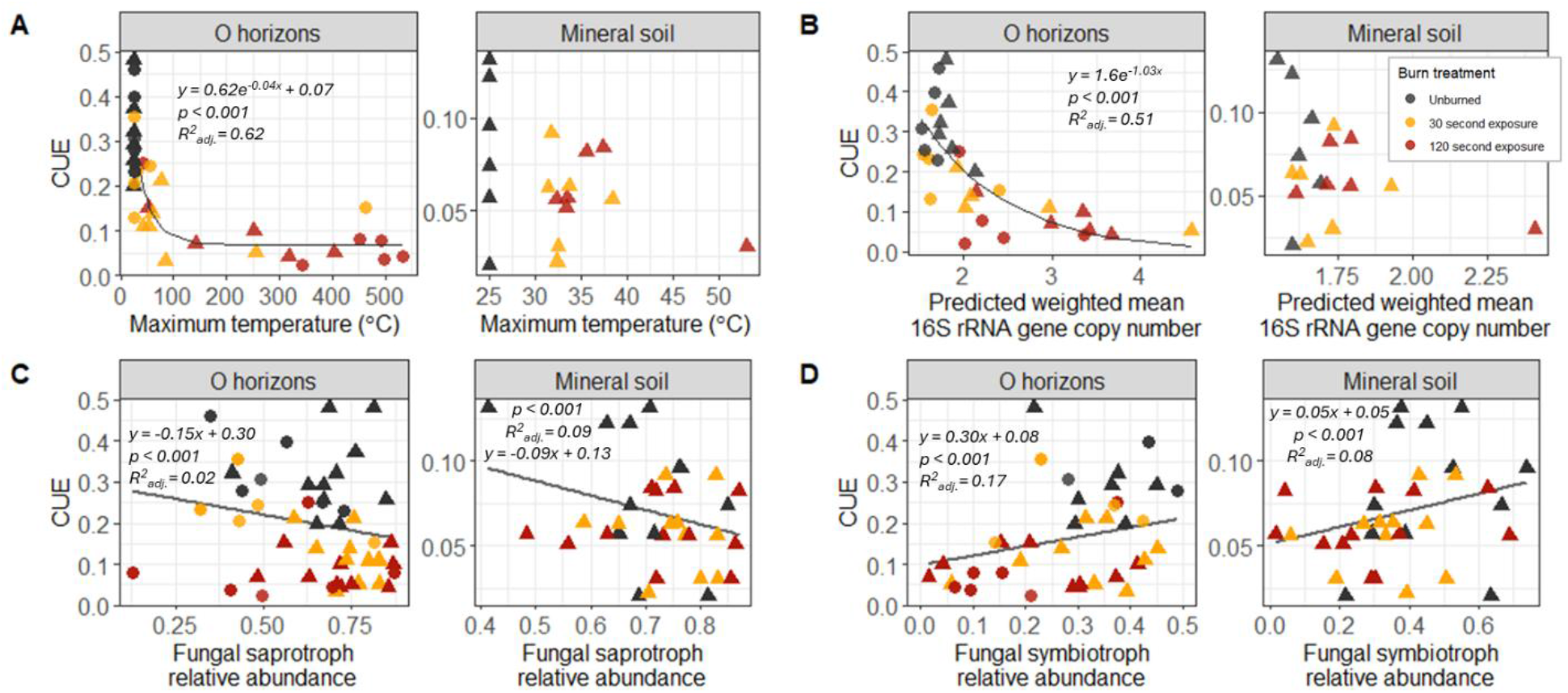
Correlation between CUE and (A) maximum temperature during burn, (B) weighted mean predicted 16S rRNA gene copy number, (C) relative abundance of putative fungal saprotrophs, and (D) relative abundance of putative fungal symbiotrophs in organic (left panels) and mineral (right panels) soil. Black, orange, and red represent soils exposed to the 0 s (unburned controls), 30 s, and 120 s burn treatment durations, respectively. Circles and triangles represent Histosols and Gleysols, respectively.

To assess the relationship between microbial community structure and function, we compared weighted mean predicted 16S rRNA gene copy numbers for bacterial communities and compared these values with measured CUE. In organic horizons, CUE decreased exponentially with increasing weighted mean predicted 16S rRNA gene copy number (Figure 6B; y = 1.6e^-1.03x^, p < 0.001, R^2^_adj._ = 0.51). In mineral horizons, there was no significant relationship between gene copy number and CUE (*p*= 0.17).

We also assessed the relationship between CUE and fungal community composition by comparing the relative abundance of putative fungal saprotrophs and symbiotrophs vs. CUE (Figure 6C & 6D). CUE decreased with increasing relative abundance of putative saprotrophs in both organic soils (y = -0.15x + 0.39, *p* = 0.05, R^2^_adj._ = 0.02) and mineral horizons (y = -0.09x + 0.05, *p* < 0.001, R^2^_adj._ = 0.09). Increasing relative abundance of putative fungal symbiotrophs correlated with increasing CUE in both organic soils (y = 0.31x + 0.08, *p* < 0.001, R^2^_adj._= 0.18) and mineral horizons (y = 0.05x + 0.05, *p* < 0.001, R^2^_adj._= 0.07). However, the explanatory power of these relationships (R^2^ ranging from 0.02 to 0.18) was much lower than that of weighted mean predicted 16S rRNA gene copy numbers (R^2^_adj._ = 0.51).

## 4. Discussion

### 4.1. Burning causes an immediate and persistent shift in microbial community composition

Laboratory burns captured realistic effects of burning on microbial community composition with greater shifts in microbial community composition following hotter burns (Figure 2), in line with previous observations of microbial communities after natural wildfires (Whitman et al., 2019; Nelson et al., 2022; Johnson et al., 2023).

Overall resistance of bacterial community composition to burning was higher than that of fungal community composition, reflecting a higher heating sensitivity of fungi than bacteria and archaea. This is unsurprising given that previous researchers have found larger burn-induced decreases in fungal vs. bacterial biomass (Pressler et al., 2019) and lower fungal:bacterial ratios in burned vs. unburned soil (Adkins et al., 2020). While there would obviously be vast variability within these different domains and kingdoms of life, this variation in resistance to burning between bacterial vs. fungal communities points towards differences in fire effects on soil organisms and suggests that increasing fire severity could differentially impact fungal communities. Although their resistance was lower, fungal communities did not show lower resilience to burning than bacterial communities in the weeks to months post-burn, suggesting that differences in bacterial vs. fungal community response to fire over time are largely driven by the initial impact of burning on microbial communities rather than differences in bacterial vs. fungal responses to post-burn soil conditions. (Figure 7)

**Figure 7.**
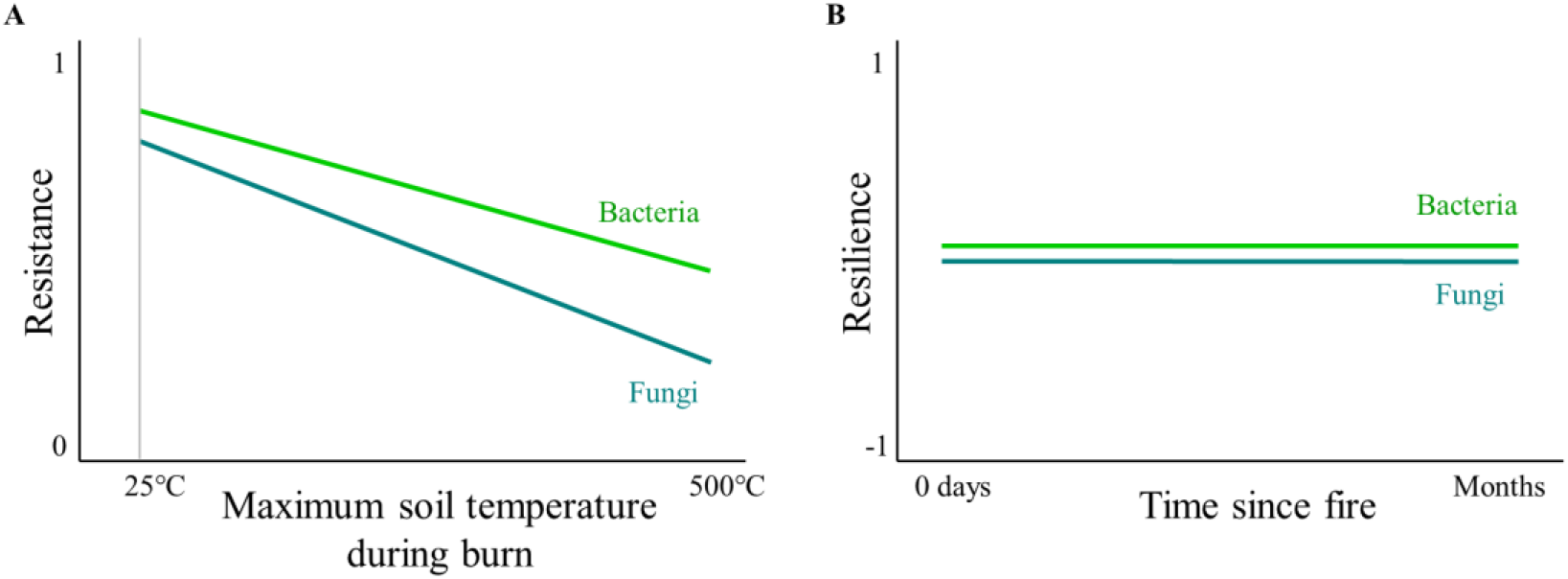
Conceptual figure showing relationships between (A) maximum burn temperature and microbial community resistance to burning and (B) time since fire and microbial community resilience to burning demonstrating similarities and differences in how bacterial (green) and fungal (blue) communities respond to fire.

Overall, low microbial community resilience to burning and the lack of trend with time since fire was counter to our predictions and suggests that either the recovery of microbial communities towards pre-burn composition occurs on longer timescales than measured in this study or that microbial communities are not on a trajectory to recover. Previous research has documented no difference in burned vs. unburned soil microbial community composition within years to decades of wildfire in boreal forest ecosystems (Treseder et al., 2004; Xiang et al., 2014), and evidence of the resilience of soil bacterial (but not fungal) communities to burning has been documented within five years following wildfire (T. Whitman et al., 2022; Whitman et al., 2025). Therefore, we speculate that microbial communities are typically resilient to fire on long timescales, and that substantial microbial community recovery occurs over years rather than weeks, with longer timescales potentially required for fungal than bacterial communities. In a study of chaparral shrubland, neither bacterial nor fungal rRNA gene abundances as measured through qPCR recovered to levels seen in unburned soils within one year of fire although bacterial 16S gene abundances did begin to increase within 6 months of the fire (Pulido-Chavez et al., 2023), further evidence of recovery occurring over months to years and of a potential difference in recovery timelines for bacteria vs. fungi. However, the laboratory burns and incubations used in our research are of course insufficient for exploring the factors shaping bacterial and fungal community recovery years to decades post-burn. Clearly, future field sampling studies will be needed to fully understand the mechanisms shaping soil bacterial and fungal recovery post-fire over decadal timescales. Based on previous literature and the lack of evidence for recovery in our study, we speculate that microbial communities in this study are on a trajectory to recover to pre-burn composition but that, in the field following natural wildfires, this recovery likely occurs over timescales longer than weeks to months and may vary across soil types and horizons.

Returning to resistance, our observations of variation in both bacterial and fungal community composition resistance to burning varied across soil horizons, with higher resistance observed in Gleysol mineral soil vs. O horizon soil (Figure 3), supporting our hypothesis that microbial community resistance to burning would be lower in O horizon vs. mineral soil. Soil temperature during a burn rapidly attenuates with depth (Beadle, 1940; Johnson et al., 2024), and, thus, microbial communities deeper in soil likely experience less heat-induced mortality and less extreme burn-induced changes in the soil environment. While the resistance of microbial communities in underlying mineral soil is high, this likely does not reflect inherent differences in the abilities of these microbial communities to withstand burning, but – rather – is the result of the dampening effects of overlying soil on burn effects. This is supported by the correlation in resistance of pH and microbial community composition (Figure 3C & D) – i.e., soils that experienced greater burn-induced shifts in pH also had larger shifts in microbial community composition, highlighting how microbial communities and characteristics of the soil environment are tightly linked, although the causality of this relationship is unclear.

We cannot disentangle whether burn-induced shifts in pH are driving changes in microbial community composition by two days post-burn, if burning is altering soil pH and microbial community composition independently, or some combination of these two explanations is playing out. Rapid shifts in microbial community composition and activity have been documented following manipulations of soil pH in laboratory studies (Brenzinger et al., 2015; Anderson et al., 2018), which suggests that the increase in soil pH observed following the burns in this study could be contributing to the shift in microbial community composition. At the same time, there are clear mechanisms by which burning directly alters microbial communities – e.g., heat-induced mortality with higher mortality following longer, hotter burns (Pingree and Kobziar, 2019; Johnson et al., 2023).

Over longer timescales of weeks to months, the link between soil pH and microbial community resistance disappears in this study, raising the question: Do the effects of burning on microbial community composition and the effects of soil pH on microbial community composition act in concert or in opposition over time? While soil pH has been shown to be a strong predictor of microbial community composition (Fierer et al., 2009; Rousk et al., 2010), the lack of correlation between pH resilience and microbial community resilience to burning suggests that the post-burn trajectory of soil pH has a minimal effect on microbial communities. Furthermore, there is no lingering effect of the initial burn-induced increase in soil pH on microbial communities over time (i.e., soil pH two days post-burn was not significantly correlated with microbial community resilience), suggesting that other factors are shaping post-burn microbial communities. The recovery of microbial communities following fire likely depends on multiple factors, including conditions in the soil environment, but also the recovery of plants and the survival of microbes available to recolonize burned soil, with the importance of these factors in structuring post-fire microbial communities varying with time since fire.

Immediately after a burn, soil microbial community composition is likely driven in large part by the combination of burn-induced mortality and effects on the soil environment, with microbial communities in deeper soil horizons being less affected by burning. This explanation is supported by both our observation that bacterial community resistance in Gleysols was higher in soil cores with thicker O horizons (Figure S9), and results from previous studies documenting larger impacts of fire on microbial communities in shallower vs. deeper soil (Caiafa et al., 2023). At first glance, our observations of higher fungal community resistance to burning in Histosols vs. Gleysol O horizons may seem to contradict this link, because burn temperatures in Histosols (5 cm below the soil surface) were higher on average than temperatures in Gleysol O horizons or mineral soil, as previously reported (Johnson et al., 2024). However, the entirety of Gleysol O horizons, which were 2.4 cm in thickness pre-burn on average, experienced higher maximum temperatures (mean = 163 °C) than the base of the Histosol soil cores, which were 10 cm thick and reached 57 °C on average during the longer 120 s burn treatments. We speculate that soil at the base of the Histosol cores may effectively serve as a fire refugium for microbes (Meddens et al., 2018), and that by homogenizing soil cores by horizon before collecting samples for DNA extraction and sequencing, we may have obscured trends in microbial community resistance with depth in Histosols. More work is needed to assess the importance of deeper soil horizons serving as microbial refugia from wildfire in the field. It is likely that following lower severity fires, microbes in unburned O horizon soils could serve as a meaningful source of microbes to colonize adjacent burned areas via dispersal by water or wind. Future work should explore whether and how quickly dispersal of soil microbes from unburned to burned areas occurs and the importance of unburned soil microbial communities in structuring microbial community composition of adjacent burn areas.

While microbial survival is obviously important for post-burn microbial community composition, microbial community composition weeks to months post-burn was not driven solely by heat-induced microbial mortality. The increase in weighted mean predicted 16S rRNA gene copy number (Figure 2) and decrease in relative abundance of fungal symbiotrophs (Figure 2) in burned soils two days post-burn indicates that fire is causing a rapid restructuring of microbial communities beyond simply killing a representative fraction of the community. The correlation between soil respiration and bacterial community resistance (Figure 3E) suggests that the burn-induced restructuring of bacteria communities may alter microbial community function in ways that impact soil C cycling immediately post-burn. Notably, respiration resistance and resilience are calculated on the basis of per g C remaining post-burn and thus reflect shifts in C utilization rather than a burn-induced decrease in total soil C. This raises the question: To what degree are post-burn changes in soil respiration driven by shifts in the size and/or composition of microbial communities? The relationship between burn-induced shifts in microbial community composition vs. function also varies with time since burn. Specifically, respiration resistance is positively correlated with bacterial resistance (R^2^_adj._ = 0.4); however, the relationship between resilience of soil respiration and resilience of bacterial communities weeks to months later is much weaker (R^2^_adj._ = 0.05). This underscores the high functional redundancy of microbial communities, where large shifts in composition do not necessarily impair ability to degrade available C substrates.

Overall, these results highlight one way in which shifts in wildfire regimes may leave lasting impacts on soil microbial communities, although the effects on composition and function may differ. If shifting wildfire regimes result in thinner O horizons due to higher severity fires (i.e., greater combustion of soil surface horizons) and less time to recover between fires (i.e., shorter fire return intervals), then we predict microbial community resistance to burning will decline, which might further reduce soil respiration rates in the days immediately post-burn. We expect that the impacts of shifting fire regimes on soil microbial communities will likely differ across soil types, and hypothesize that any decrease in microbial community resistance to burning under future wildfire regimes would be concentrated in the O horizon.

### 4.2. C cycling in post-burn soils may be affected by shift in bacterial communities towards faster growth and lower efficiency

Our observations of decreased microbial community CUE 24 days post-burn in burned vs. unburned organic horizon soils from both soil types (Figure 5C) were consistent with our hypothesis that burning would reduce glucose-specific CUE. The observed decrease in CUE post-burn was driven primarily by decreased microbial growth – burning had a small positive effect on glucose-derived respiration, which is evidence that the microbial community in burned soils is still active and degrading the added glucose 24 days post-burn. At the same time, we observed substantially lower incorporation of C from glucose into microbial biomass (i.e., microbial growth) in burned soils, hence the decrease in CUE in burned organic horizon soils.

We posit that decreased CUE is caused in part by an increase in the relative abundance of fast-growing bacterial taxa, which may be driven by an increase in available C caused by the death of microbes and plant roots during the burn. The heat-induced production of necromass during a fire may stimulate an increase in fast-growing microbial taxa dependent upon simple C substrates that are easily accessible to microbes (i.e., not sorbed to mineral surfaces). A shift in C availability towards more accessible C substrates is consistent with the observed decrease in glucose-specific CUE. While measurements of glucose-specific CUE reflect shifts in microbial community utilization of the added glucose and do not directly reflect burn-induced changes to soil C chemistry (i.e., necromass or pyrogenic organic matter production), shifts in soil C chemistry may indirectly affect glucose-specific CUE via changes in microbial community composition. This highlights how burn-induced shifts in soil C pools may affect microbially medicated C cycling for months post-burn.

Previous field research in wildfires in this region has demonstrated an increase in the relative abundance of fast-growing bacterial taxa following fire, which was positively correlated with burn severity and weighted mean predicted 16S rRNA gene copy number (Johnson et al., 2023).

This was supported by our observations of an exponential decrease in CUE with increasing weighted mean predicted 16S rRNA gene copy number (Figure 6B), which has been shown to correlate with bacterial maximum potential growth rates (Roller et al., 2016). Thus, while putatively fast-growing bacterial taxa in burned soils seem to be mineralizing glucose in burned soils at a rate similar to or faster than unburned microbial communities, these fast growers appear to be less efficient at incorporating C from glucose into biomass.

One possible explanation for our observations of CUE is a growth rate-yield tradeoff, i.e., increased microbial investment in resource acquisition to fuel faster growth comes at the cost of efficiency in the form of increased respiration and futile metabolic cycles necessary to offset stoichiometric imbalances within the cell (Manzoni et al., 2018). An alternative explanation is that burn-induced shifts in the soil environment may prompt a change in C allocation strategies for microbes due to increased environmental stress. This could manifest as an increase in maintenance respiration, leading to decreased CUE (Manzoni et al., 2018) and/or an increase in C allocation to extracellular products such as extracellular enzymes or exopolysaccharides. Data on microbial exudates in the weeks to months post-fire is sparse but suggests fire effects on extracellular products likely vary substantially with time since fire (Goberna et al., 2012; Dove et al., 2020; Pellegrini et al., 2021; Szafrańska et al., 2026). Future laboratory work exploring the impact of burning on microbial maintenance respiration and exudate concentrations days to weeks post-burn is needed to disentangle the microbial mechanisms driving decreased CUE post-burn. Even in the absence of a mechanistic understanding of the relationship between burning and CUE, we can still begin to explore how shifting wildfire regimes may impact post-burn microbial community CUE. The negative relationship between CUE and maximum soil burn temperatures suggests that the higher energy absorption from hotter and/or slower moving fires will depress CUE more, and thus, have larger impacts on soil C cycling in the months to years following fire. These changes in CUE could have important implications for C stocks. If microbes process C differently after a burn by incorporating proportionally less C into biomass, that would also result in less necromass when the microbes die. Because of the close interactions between living microbes and mineral surfaces, their necromass may disproportionately contribute to mineral-associated organic matter, which is thought to be more persistent than the soil C precursors to microbial biomass (Cotrufo et al., 2013). Thus, determining the timescale over which this effect on CUE persists will be important in understanding the scale of any potential downstream effects on soil C stocks.

Previous work has shown that the enrichment of fast-growing taxa can persist for at least one year following fire in both organic and mineral soils from this region (Johnson et al., 2023). If the observed decreases in CUE in organic soil following burning are driven primarily by a shift in microbial community composition to more fast-growing, less-efficient taxa, then it is possible that the return of CUE to pre-burn levels could require a year or more. However, our measurements of CUE integrate the activity of the entire microbial community, including bacteria, archaea, and fungi, whereas 16S rRNA gene copy number captures only bacteria and archaea. Thus, the recovery of CUE to pre-burn levels may not mirror exactly the dynamics of the bacterial community. While fast-growing fungal taxa may also thrive in the post-burn soil environment, there is not currently a robust community composition-based way to predict relative growth rates of fungal communities. Burn-induced shifts in fungal trophic strategies may be contributing to decreased microbial community CUE as suggested by the correlations between CUE and the relative abundances of putative saprotrophic and symbiotrophic fungi (Figure 6).

However, these correlations between fungal communities and CUE are much weaker than the relationship between predicted weighted mean 16S rRNA gene copy number and CUE, and evidence for causal impacts of fungal community composition on CUE is lacking. Clearly more work is needed to understand how burn-induced shifts in fungal community composition relate to shifts in CUE.

Our CUE data are temporally limited, representing a single timepoint of substrate-specific CUE post-burn, and there are numerous ways in which the effects of fire on glucose-specific CUE, as well as CUE of all soil C substrates, may vary over time. The combination of elevated 16S rRNA gene copy numbers in burned soils from 2 to 70 days post-burn (Figure 2B) and the strong negative correlation between gene copy numbers and glucose-specific CUE at 24 days post-burn (Figure 6B) prompts us to speculate that burning may cause a rapid and persistent decrease in CUE, but clearly more work is needed to experimentally test the relationship between gene copy number and CUE in burned soils over long timescales.

The observed relatively strong negative correlation (R^2^ = 0.51) between CUE and weighted mean predicted 16S rRNA gene copy number was also intriguing to us because of its potentially important implications for modelling. Many soil C cycling models rely on CUE as a central parameter (Allison, 2025), and recent work indicates that CUE is critically important in driving global models of SOC stocks (Tao et al., 2023). Given the relationship between CUE, weighted mean predicted 16S rRNA gene copy number, and fast-growing taxa, the use of sequencing data to potentially inform the partitioning of fast-growing/low-CUE and slow-growing/high-CUE soil microbial pools in microbially-explicit models should be explored. However, the broad existence and potential utility of this correlation currently remains highly speculative – the next step would be to assess the existence of this relationship across more substrates, communities, and conditions.

While there are numerous ways in which the effects of fire on microbial community recovery, growth rates, and CUE may vary over time – e.g., changes in soil temperature, shifts in plant inputs – that are not explored here, these findings increase our understanding of microbial dynamics immediately post-burn and open questions about the timeline and trajectory of the recovery of microbial community composition and function following fire. Shifting wildfire regimes towards hotter fires and/or towards shorter fire return intervals possibly leading to shallower organic horizons could impact C cycling via larger and more persistent changes in microbial community composition with concomitant changes in CUE. Larger decreases in CUE would impact soil C stocks via a higher proportion of decomposed soil organic matter being transported to the atmosphere as CO_2_.

## 5. Conclusion

Burning causes an immediate but nuanced shift in microbial community composition with bacterial and fungal community resistance to burning varying across soil type. The relatively low resilience of bacterial and fungal communities to burning as well as the failure of resilience to increase with time since burning is evidence that timescales longer than 10 weeks are needed to observe the beginning of microbial community recovery towards pre-burn composition. In organic horizons, burning also caused a decrease in CUE, which correlated with an exponential increase in weighted mean predicted 16S rRNA gene copy numbers, supporting our hypothesis that burning causes a shift towards faster growing, less efficient microbial communities. The effects of burning on CUE were larger following burns of higher maximum soil temperature and minimal in underlying mineral soil, which showcases the nuance of burn-induced changes in soil C cycling and highlights ways in which the effects of shifting boreal forest wildfire regimes may vary across soil types and depths. Together, these results support previous calls for a more nuanced representation of CUE in soil C models (Manzoni et al., 2018; He et al., 2024; Allison, 2025).

## Acknowledgements

The authors thank the University of Wisconsin Biotechnology Center DNA Sequencing Facility (Research Resource Identifier – RRID:SCR_017759) for providing Illumina sequencing facilities and services.

We thank D. Letourneau and M.A. Parisien at Natural Resources Canada for assistance with field work; J. Morin, T.J. Little, and other Wood Buffalo National Park staff for support in conducting this research (Permit WB-2022-41998); E. Lazarcik and K. Bourne at the USDA FS Forest Products Laboratory for facilitating the burn simulations; Z. Freedman for support with headspace gas CO_2_-C analysis; T. Berry for help with ^13^C isotope measurements and calibrations; K. Kruger and A. Hutka for assistance with laboratory work; N. Zeba, M. Sikora, and J. Woolet for producing the ^13^C-labelled pine material; H. Read for assistance with data acquisition; and anonymous reviewers for their helpful comments on the manuscript.

## Author contribution statement

**Dana Johnson**: Conceptualization, Methodology, Formal analysis, Investigation, Data curation, Writing – original draft, Visualization. **Kara Yedinak**: Methodology, Writing – review and editing. **Thea Whitman**: Funding acquisition, Conceptualization, Methodology, Resources, Writing – review and editing.

No generative artificial intelligence tools were used for any aspect of this paper, including generating the ideas, analyzing the data, or writing the manuscript.

## Data availability statement

The sequencing datasets generated during the current study are available in the NCBI SRA under bioproject number PRJNA1369477. Non-sequencing data are available in the DOE ESS-DIVE repository at https://doi.org/10.15485/2438579. Source data are provided in this paper.

Code to analyze data and produce figures is available at GitHub at https://github.com/DanaBJohnson/Microbial-Resistance-and-Resilience

## References

Abarenkov, K., Nilsson, R.H., Larsson, K.-H., Taylor, A.F.S., May, T.W., Frøslev, T.G., Pawlowska, J., Lindahl, B., Põldmaa, K., Truong, C., Vu, D., Hosoya, T., Niskanen, T., Piirmann, T., Ivanov, F., Zirk, A., Peterson, M., Cheeke, T.E., Ishigami, Y., Jansson, A.T., Jeppesen, T.S., Kristiansson, E., Mikryukov, V., Miller, J.T., Oono, R., Ossandon, F.J., Paupério, J., Saar, I., Schigel, D., Suija, A., Tedersoo, L., Kõljalg, U., 2024. The UNITE database for molecular identification and taxonomic communication of fungi and other eukaryotes: sequences, taxa and classifications reconsidered. Nucleic Acids Research 52, D791–D797. doi:10.1093/nar/gkad1039

Adkins, J., Docherty, K.M., Gutknecht, J.L.M., Miesel, J.R., 2020. How do soil microbial communities respond to fire in the intermediate term? Investigating direct and indirect effects associated with fire occurrence and burn severity. The Science of the Total Environment 745. doi:10.1016/J.SCITOTENV.2020.140957

Allison, S.D., 2025. Rethinking microbial carbon use efficiency in soil models. Nature Climate Change 15, 10–12. doi:10.1038/s41558-024-02217-6

Anderson, C.R., Peterson, M.E., Frampton, R.A., Bulman, S.R., Keenan, S., Curtin, D., 2018. Rapid increases in soil pH solubilise organic matter, dramatically increase denitrification potential and strongly stimulate microorganisms from the Firmicutes phylum. PeerJ 6, e6090. doi:10.7717/peerj.6090

Auwal, M., Sun, H., Adamu, U.K., Meng, J., Van Zwieten, L., Pal Singh, B., Luo, Y., Xu, J., 2023. The phosphorus limitation in the post-fire forest soils increases soil CO2 emission via declining cellular carbon use efficiency and increasing extracellular phosphatase. CATENA 224, 106968. doi:10.1016/j.catena.2023.106968

Bárcenas-Moreno, G., García-Orenes, F., Mataix-Solera, J., Mataix-Beneyto, J., Bååth, E., 2011. Soil microbial recolonisation after a fire in a Mediterranean forest. Biology and Fertility of Soils 47, 261–272. doi:10.1007/s00374-010-0532-2

Beadle, N.C.W., 1940. Soil temperatures during forest fires and their effect on the survival of vegetation. The Journal of Ecology 28, 180–192. doi:10.2307/2256168

Bodí, M.B., Martin, D.A., Balfour, V.N., Santín, C., Doerr, S.H., Pereira, P., Cerdà, A., Mataix-Solera, J., 2014. Wildland fire ash: Production, composition and eco-hydro-geomorphic effects. Earth-Science Reviews 130, 103–127.

Bokulich, N.A., Kaehler, B.D., Rideout, J.R., Dillon, M., Bolyen, E., Knight, R., Huttley, G.A., Gregory Caporaso, J., 2018. Optimizing taxonomic classification of marker-gene amplicon sequences with QIIME 2’s q2-feature-classifier plugin. Microbiome 6, 90. doi:10.1186/s40168-018-0470-z

Bolyen, E., Rideout, J.R., Dillon, M.R., Bokulich, N.A., Abnet, C.C., Al-Ghalith, G.A., Alexander, H., Alm, E.J., Arumugam, M., Asnicar, F., Bai, Y., Bisanz, J.E., Bittinger, K., Brejnrod, A., Brislawn, C.J., Brown, C.T., Callahan, B.J., Caraballo-Rodríguez, A.M., Chase, J., Cope, E.K., Da Silva, R., Diener, C., Dorrestein, P.C., Douglas, G.M., Durall, D.M., Duvallet, C., Edwardson, C.F., Ernst, M., Estaki, M., Fouquier, J., Gauglitz, J.M., Gibbons, S.M., Gibson, D.L., Gonzalez, A., Gorlick, K., Guo, J., Hillmann, B., Holmes, S., Holste, H., Huttenhower, C., Huttley, G.A., Janssen, S., Jarmusch, A.K., Jiang, L., Kaehler, B.D., Kang, K.B., Keefe, C.R., Keim, P., Kelley, S.T., Knights, D., Koester, I., Kosciolek, T., Kreps, J., Langille, M.G.I., Lee, J., Ley, R., Liu, Y.-X., Loftfield, E., Lozupone, C., Maher, M., Marotz, C., Martin, B.D., McDonald, D., McIver, L.J., Melnik, A.V., Metcalf, J.L., Morgan, S.C., Morton, J.T., Naimey, A.T., Navas-Molina, J.A., Nothias, L.F., Orchanian, S.B., Pearson, T., Peoples, S.L., Petras, D., Preuss, M.L., Pruesse, E., Rasmussen, L.B., Rivers, A., Robeson, M.S., Rosenthal, P., Segata, N., Shaffer, M., Shiffer, A., Sinha, R., Song, S.J., Spear, J.R., Swafford, A.D., Thompson, L.R., Torres, P.J., Trinh, P., Tripathi, A., Turnbaugh, P.J., Ul-Hasan, S., van der Hooft, J.J.J., Vargas, F., Vázquez-Baeza, Y., Vogtmann, E., von Hippel, M., Walters, W., Wan, Y., Wang, M., Warren, J., Weber, K.C., Williamson, C.H.D., Willis, A.D., Xu, Z.Z., Zaneveld, J.R., Zhang, Y., Zhu, Q., Knight, R., Caporaso, J.G., 2019. Reproducible, interactive, scalable and extensible microbiome data science using QIIME 2. Nature Biotechnology 37, 852–857. doi:10.1038/s41587-019-0209-9

Borgogni, F., Lavecchia, A., Mastrolonardo, G., Certini, G., Ceccherini, M.T., Pietramellara, G., 2019. Immediate- and Short-term Wildfire Impact on Soil Microbial Diversity and Activity in a Mediterranean Forest Soil. Soil Science 184, 35. doi:10.1097/SS.0000000000000250

Bradshaw, C.J.A., Warkentin, I.G., 2015. Global estimates of boreal forest carbon stocks and flux. Global and Planetary Change 128, 24–30. doi:10.1016/j.gloplacha.2015.02.004

Bradshaw, M.J., Aime, M.C., Rokas, A., Maust, A., Moparthi, S., Jellings, K., Pane, A.M., Hendricks, D., Pandey, B., Li, Y., Pfister, D.H., 2023. Extensive intragenomic variation in the internal transcribed spacer region of fungi. iScience 26, 107317. doi:10.1016/j.isci.2023.107317

Bray, J.R., Curtis, J.T., 1957. An Ordination of the Upland Forest Communities of Southern Wisconsin. Ecological Monographs 27, 325–349. doi:10.2307/1942268

Brenzinger, K., Dörsch, P., Braker, G., 2015. pH-driven shifts in overall and transcriptionally active denitrifiers control gaseous product stoichiometry in growth experiments with extracted bacteria from soil. Frontiers in Microbiology 6. doi:10.3389/fmicb.2015.00961

Caiafa, M.V., Nelson, A.R., Borch, T., Roth, H.K., Fegel, T.S., Rhoades, C.C., Wilkins, M.J., Glassman, S.I., 2023. Distinct fungal and bacterial responses to fire severity and soil depth across a ten-year wildfire chronosequence in beetle-killed lodgepole pine forests. Forest Ecology and Management 544, 121160. doi:10.1016/J.FORECO.2023.121160

Callahan, B.J., McMurdie, P.J., Rosen, M.J., Han, A.W., Johnson, A.J.A., Holmes, S.P., 2016. DADA2: High-resolution sample inference from Illumina amplicon data. Nature Methods 13, 581–583. doi:10.1038/nmeth.3869

Certini, G., 2005. Effects of fire on properties of forest soils: A review. Oecologia 143, 1–10. doi:10.1007/s00442-004-1788-8

Cobo-Díaz, J.F., Fernández-González, A.J., Villadas, P.J., Robles, A.B., Toro, N., Fernández-López, M., 2015. Metagenomic Assessment of the Potential Microbial Nitrogen Pathways in the Rhizosphere of a Mediterranean Forest After a Wildfire. Microbial Ecology 69, 895–904. doi:10.1007/s00248-015-0586-7

Cotrufo, M.F., Wallenstein, M.D., Boot, C.M., Denef, K., Paul, E., 2013. The Microbial Efficiency-Matrix Stabilization (MEMS) framework integrates plant litter decomposition with soil organic matter stabilization: do labile plant inputs form stable soil organic matter? Global Change Biology 19, 988–995.

de Groot, W.J., Cantin, A.S., Flannigan, M.D., Soja, A.J., Gowman, L.M., Newbery, A., 2013a. A comparison of Canadian and Russian boreal forest fire regimes. Forest Ecology and Management, The Mega-fire reality 294, 23–34. doi:10.1016/j.foreco.2012.07.033

de Groot, W.J., Flannigan, M.D., Cantin, A.S., 2013b. Climate change impacts on future boreal fire regimes. Forest Ecology and Management 294, 35–44. doi:10.1016/j.foreco.2012.09.027

DeBano, L.F., 1991. The effect of fire on soil properties, in: Harvey, A.E., Neuenschwander, L.F. (Eds.), Proceedings-Management and Productivity of Western-Montane Forest Soils; 1990 April 10-12; Boise, ID. Gen. Tech. Rep. INT-280. Ogden, UT: U.S. Department of Agriculture, Forest Service, Intermountain Research Station, Berkeley, California, USA, pp. 151–156.

Dictor, M.-C., Tessier, L., Soulas, G., 1998. Reassessement of the *K*ec coefficient of the fumigation–extraction method in a soil profile. Soil Biology and Biochemistry 30, 119–127. doi:10.1016/S0038-0717(97)00111-9

Dove, N.C., Safford, H.D., Bohlman, G.N., Estes, B.L., Hart, S.C., 2020. High-severity wildfire leads to multi-decadal impacts on soil biogeochemistry in mixed-conifer forests. Ecological Applications 30, e02072. doi:10.1002/EAP.2072

Dove, N.C., Taş, N., Hart, S.C., 2022. Ecological and genomic responses of soil microbiomes to high-severity wildfire: linking community assembly to functional potential. The ISME Journal 16, 1853–1863. doi:10.1038/s41396-022-01232-9

Fierer, N., Strickland, M.S., Liptzin, D., Bradford, M.A., Cleveland, C.C., 2009. Global patterns in belowground communities. Ecology Letters 12, 1238–1249. doi:10.1111/j.1461-0248.2009.01360.x

Food and Agriculture Organization of the United Nations., 2003. Digital soil map of the world and derived soil properties [WWW Document]. Land and Water Development Division. Unesco. URL (accessed 3.22.22).

Frankman, D., Webb, B.W., Butler, B.W., Jimenez, D., Forthofer, J.M., Sopko, P., Shannon, K.S., Hiers, J.K., Ottmar, R.D., 2012. Measurements of convective and radiative heating in wildland fires. International Journal of Wildland Fire 22, 157–167. doi:10.1071/WF11097

Geyer, K.M., Dijkstra, P., Sinsabaugh, R., Frey, S.D., 2019. Clarifying the interpretation of carbon use efficiency in soil through methods comparison. Soil Biology and Biochemistry 128, 79–88. doi:10.1016/J.SOILBIO.2018.09.036

Geyer, K.M., Kyker-Snowman, E., Grandy, A.S., Frey, S.D., 2016. Microbial carbon use efficiency: accounting for population, community, and ecosystem-scale controls over the fate of metabolized organic matter. Biogeochemistry 127, 173–188. doi:10.1007/s10533-016-0191-y

Glassman, S.I., Levine, C.R., Dirocco, A.M., Battles, J.J., Bruns, T.D., 2016. Ectomycorrhizal fungal spore bank recovery after a severe forest fire: Some like it hot. ISME Journal 10, 1228–1239. doi:10.1038/ismej.2015.182

Global Wildfire Information System, 2026. “Annual area burnt by wildfires” [dataset].

Glöckner, F.O., Yilmaz, P., Quast, C., Gerken, J., Beccati, A., Ciuprina, A., Bruns, G., Yarza, P., Peplies, J., Westram, R., Ludwig, W., 2017. 25 years of serving the community with ribosomal RNA gene reference databases and tools. Journal of Biotechnology 261, 169–176. 10.1016/j.jbiotec.2017.06.1198

Goberna, M., García, C., Insam, H., Hernández, M.T., Verdú, M., 2012. Burning Fire-Prone Mediterranean Shrublands: Immediate Changes in Soil Microbial Community Structure and Ecosystem Functions. Microbial Ecology 64, 242–255. doi:10.1007/s00248-011-9995-4

He, X., Abramoff, R.Z., Abs, E., Georgiou, K., Zhang, H., Goll, D.S., 2024. Model uncertainty obscures major driver of soil carbon. Nature 627, E1–E3. doi:10.1038/s41586-023-06999-1

Holden, S.R., Rogers, B.M., Treseder, K.K., Randerson, J.T., 2016. Fire severity influences the response of soil microbes to a boreal forest fire. Environmental Research Letters 11, 1–10. doi:10.1088/1748-9326/11/3/035004

Johnson, D.B., Woolet, J., Yedinak, K.M., Whitman, T., 2023. Experimentally determined traits shape bacterial community composition one and five years following wildfire. Nature Ecology & Evolution 7, 1419–1431. doi:10.1038/s41559-023-02135-4

Johnson, D.B., Yedinak, K.M., Sulman, B.N., Berry, T.D., Kruger, K., Whitman, T., 2024. Effects of fire and fire-induced changes in soil properties on post-burn soil respiration. Fire Ecology 20, 90. 10.1186/s42408-024-00328-1

Kasischke, E.S., French, N.H.F., Bourgeau-Chavez, L.L., Christensen Jr., N.L., 1995. Estimating release of carbon from 1990 and 1991 forest fires in Alaska. Journal of Geophysical Research: Atmospheres 100, 2941–2951. doi:10.1029/94JD02957

Kasischke, E.S., Johnstone, J.F., 2005. Variation in postfire organic layer thickness in a black spruce forest complex in interior Alaska and its effects on soil temperature and moisture. Canadian Journal of Forest Research 35, 2164–2177. doi:10.1139/x05-159

Kasischke, E.S., Turetsky, M.R., 2006. Recent changes in the fire regime across the North American boreal region—Spatial and temporal patterns of burning across Canada and Alaska. Geophysical Research Letters 33, L09703. doi:10.1029/2006GL025677

Kasischke, E.S., Verbyla, D.L., Rupp, T.S., McGuire, A.D., Murphy, K.A., Jandt, R., Barnes, J.L., Hoy, E.E., Duffy, P.A., Calef, M., Turetsky, M.R., 2010. Alaska’s changing fire regime--implications for the vulnerability of its boreal forests. Canadian Journal of Forest Research. 40: 1313–1324 40, 1313–1324.

Kelly, J., Ibáñez, T.S., Santín, C., Doerr, S.H., Nilsson, M.C., Holst, T., Lindroth, A., Kljun, N., 2021. Boreal forest soil carbon fluxes one year after a wildfire: Effects of burn severity and management. Global Change Biology 27, 4181–4195. doi:10.1111/GCB.15721

Kelly, R., Chipman, M.L., Higuera, P.E., Stefanova, I., Brubaker, L.B., Hu, F.S., 2013. Recent burning of boreal forests exceeds fire regime limits of the past 10,000 years. Proceedings of the National Academy of Sciences of the United States of America 110, 13055–13060. doi:10.1073/PNAS.1305069110/SUPPL_FILE/ST01.DOC

Kozich, J.J., Westcott, S.L., Baxter, N.T., Highlander, S.K., Schloss, P.D., 2013. Development of a dual-index sequencing strategy and curation pipeline for analyzing amplicon sequence data on the MiSeq Illumina sequencing platform. Applied and Environmental Microbiology 79, 5112–5120. doi:10.1128/AEM.01043-13

Lauber, C.L., Hamady, M., Knight, R., Fierer, N., 2009. Pyrosequencing-based assessment of soil pH as a predictor of soil bacterial community structure at the continental scale. Applied and Environmental Microbiology 75, 5111–5120. doi:10.1128/AEM.00335-09

Lofgren, L.A., Uehling, J.K., Branco, S., Bruns, T.D., Martin, F., Kennedy, P.G., 2019. Genome-based estimates of fungal rDNA copy number variation across phylogenetic scales and ecological lifestyles. Molecular Ecology 28, 721–730. doi:10.1111/mec.14995

Lou, H., Cai, H., Fu, R., Guo, C., Fan, B., Hu, H., Zhang, J., Sun, L., 2023. Effects of wildfire disturbance on forest soil microbes and colonization of ericoid mycorrhizal fungi in northern China. Environmental Research 231, 116220. doi:10.1016/J.ENVRES.2023.116220

Luo, M., 2023. Impacts of subsequent fire on physical, chemical, and biological characteristics of preexisting pyrogenic organic matter. Dissertation. University of Wisconsin-Madison, Madison, Wisconsin, USA.

Makarov, M.I., Malysheva, T.I., Menyailo, O.V., Soudzilovskaia, N.A., Van Logtestijn, R.S.P., Cornelissen, J.H.C., 2015. Effect of KSO concentration on extractability and isotope signature (δC and δN) of soil C and N fractions. European Journal of Soil Science 66, 417–426. 10.1111/ejss.12243

Manzoni, S., Capek, P., Porada, P., Thurner, M., Winterdahl, M., Beer, C., Brüchert, V., Frouz, J., Herrmann, A.M., Lindahl, B.D., Lyon, S.W., Santrucková, H., Vico, G., Way, D., 2018. Reviews and syntheses: Carbon use efficiency from organisms to ecosystems - definitions, theories, and empirical evidence. BIOGEOSCIENCES 15, 5929–5949. doi:10.5194/bg-15-5929-2018

Martin, M., 2011. Cutadapt removes adapter sequences from high-throughput sequencing reads. EMBnet.Journal 17, 10–12. doi:10.14806/ej.17.1.200

Masyagina, O.V., Tokareva, I.V., Prokushkin, A.S., 2016. Post fire organic matter biodegradation in permafrost soils: Case study after experimental heating of mineral horizons. Science of The Total Environment 573, 1255–1264. doi:10.1016/j.scitotenv.2016.04.195

McMurdie, P.J., Holmes, S., 2013. Phyloseq: An R package for reproducible interactive analysis and graphics of microbiome census data. PLoS ONE 8, e61217. doi:10.1371/journal.pone.0061217

Meddens, A.J.H., Kolden, C.A., Lutz, J.A., Smith, A.M.S., Cansler, C.A., Abatzoglou, J.T., Meigs, G.W., Downing, W.M., Krawchuk, M.A., 2018. Fire Refugia: What Are They, and Why Do They Matter for Global Change? BioScience 68, 944–954. doi:10.1093/biosci/biy103

Neary, D.G., Klopatek, C.C., DeBano, L.F., Ffolliott, P.F., 1999. Fire effects on belowground sustainability: A review and synthesis. Forest Ecology and Management 122, 51–71. doi:10.1016/S0378-1127(99)00032-8

Nelson, A.R., Narrowe, A.B., Rhoades, C.C., Fegel, T.S., Daly, R.A., Roth, H.K., Chu, R.K., Amundson, K.K., Young, R.B., Steindorff, A.S., Mondo, S.J., Grigoriev, I.V., Salamov, A., Borch, T., Wilkins, M.J., 2022. Wildfire-dependent changes in soil microbiome diversity and function. Nature Microbiology 2022 7:9 7, 1419–1430. doi:10.1038/s41564-022-01203-y

Nelson, A.R., Rhoades, C.C., Fegel, T.S., Roth, H.K., Caiafa, M.V., Glassman, S.I., Borch, T., Wilkins, M.J., 2024. Wildfire impact on soil microbiome life history traits and roles in ecosystem carbon cycling. ISME Communications 4, ycae108. doi:10.1093/ismeco/ycae108

Nemergut, D.R., Knelman, J.E., Ferrenberg, S., Bilinski, T., Melbourne, B., Jiang, L., Violle, C., Darcy, J.L., Prest, T., Schmidt, S.K., Townsend, A.R., 2015. Decreases in average bacterial community rRNA operon copy number during succession. The ISME Journal 10, 1147–1156. doi:10.1038/ismej.2015.191

Nguyen, N.H., Song, Z., Bates, S.T., Branco, S., Tedersoo, L., Menke, J., Schilling, J.S., Kennedy, P.G., 2016. FUNGuild: An open annotation tool for parsing fungal community datasets by ecological guild. Fungal Ecology 20, 241–248. doi:10.1016/j.funeco.2015.06.006

Oksanen, J., Blanchet, F.G., Friendly, M., Kindt, R., Legendre, P., McGlinn, D., Minchin, P.R., O’Hara, R.B., Simpson, G.L., Solymos, P., Stevens, M.H.H., Szoecs, E., Wagner, H., 2020. vegan: Community Ecology Package. R Package Version 2.5–7.

Orwin, K.H., Wardle, D.A., 2004. New indices for quantifying the resistance and resilience of soil biota to exogenous disturbances. Soil Biology and Biochemistry 36, 1907–1912. 10.1016/j.soilbio.2004.04.036

Pang, R., Xu, X., Tian, Y., Cui, X., Ouyang, H., Kuzyakov, Y., 2021. In-situ 13CO2 labeling to trace carbon fluxes in plant-soil-microorganism systems: Review and methodological guideline. Rhizosphere 20, 100441. 10.1016/j.rhisph.2021.100441

Pellegrini, A.F.A., Caprio, A.C., Georgiou, K., Finnegan, C., Hobbie, S.E., Hatten, J.A., Jackson, R.B., 2021. Low-intensity frequent fires in coniferous forests transform soil organic matter in ways that may offset ecosystem carbon losses. Global Change Biology 27, 3810–3823. doi:10.1111/gcb.15648

Pérez-Valera, E., Goberna, M., Verdú, M., 2019. Fire modulates ecosystem functioning through the phylogenetic structure of soil bacterial communities. Soil Biology and Biochemistry 129, 80–89. doi:10.1016/j.soilbio.2018.11.007

Pingree, M.R.A., Kobziar, L.N., 2019. The myth of the biological threshold: A review of biological responses to soil heating associated with wildland fire. Forest Ecology and Management 432, 1022–1029. doi:10.1016/j.foreco.2018.10.032

Potthoff, M., Loftfield, N., Buegger, F., Wick, B., John, B., Joergensen, R.G., Flessa, H., 2003. The determination of δ13C in soil microbial biomass using fumigation-extraction. Soil Biology and Biochemistry 35, 947–954. 10.1016/S0038-0717(03)00151-2

Pressler, Y., Moore, J.C., Cotrufo, M.F., 2019. Belowground community responses to fire: Meta-analysis reveals contrasting responses of soil microorganisms and mesofauna. Oikos 128, 309–327. doi:10.1111/OIK.05738

Prince, T.J., Pisaric, M.F.J., Turner, K.W., 2018. Postglacial reconstruction of fire history using sedimentary charcoal and pollen from a small lake in Southwest Yukon Territory, Canada. Frontiers in Ecology and Evolution 6, 209.

Pulido-Chavez, M.F., Randolph, J.W.J.J., Zalman, C., Larios, L., Homyak, P.M., Glassman, S.I., 2023. Rapid bacterial and fungal successional dynamics in first year after chaparral wildfire. Molecular Ecology 32, 1685–1707. 10.1111/mec.16835

Quast, C., Pruesse, E., Yilmaz, P., Gerken, J., Schweer, T., Yarza, P., Peplies, J., Glöckner, F.O., 2013. The SILVA ribosomal RNA gene database project: improved data processing and web-based tools. Nucleic Acids Research 41, D590–D596. doi:10.1093/nar/gks1219

R Core Team, 2024. R A Language and Environment for Statistical Computing.

Ribeiro-Kumara, C., Santín, C., Doerr, S.H., Pumpanen, J., Baxter, G., Köster, K., 2022. Short- to medium-term effects of crown and surface fires on soil respiration in a Canadian boreal forest. Canadian Journal of Forest Research 52, 591–604. doi:10.1139/cjfr-2021-0354

Roller, B.R.K., Stoddard, S.F., Schmidt, T.M., 2016. Exploiting rRNA operon copy number to investigate bacterial reproductive strategies. Nature Microbiology 1. doi:10.1038/nmicrobiol.2016.160

Rousk, J., Bååth, E., Brookes, P.C., Lauber, C.L., Lozupone, C., Caporaso, J.G., Knight, R., Fierer, N., 2010. Soil bacterial and fungal communities across a pH gradient in an arable soil. ISME Journal 4, 1340–1351. doi:10.1038/ismej.2010.58

Scharlemann, J.P.W., Tanner, E.V.J., Hiederer, R., Kapos, V., 2014. Global soil carbon: understanding and managing the largest terrestrial carbon pool. Carbon Management 5, 81–91. doi:10.4155/cmt.13.77

Sorensen, J.W., Shade, A., 2020. Dormancy dynamics and dispersal contribute to soil microbiome resilience. Philosophical Transactions of the Royal Society B: Biological Sciences 375. doi:10.1098/rstb.2019.0255

Stocks, B.J., Lee, B.S., Martell, D.L., 1996. Some potential carbon budget implications of fire management in the boreal forest, in: Apps, M.J., Price, D.T. (Eds.), Forest Ecosystems, Forest Management and the Global Carbon Cycle. Springer Berlin Heidelberg, Berlin, Heidelberg, pp. 89–96.

Stoddard, S.F., Smith, B.J., Hein, R., Roller, B.R.K., Schmidt, T.M., 2015. rrnDB: Improved tools for interpreting rRNA gene abundance in bacteria and archaea and a new foundation for future development. Nucleic Acids Research 43, D593–D598. doi:10.1093/nar/gku1201

Sulman, B.N., Moore, J.A.M., Abramoff, R., Averill, C., Kivlin, S., Georgiou, K., Sridhar, B., Hartman, M.D., Wang, G., Wieder, W.R., Bradford, M.A., Luo, Y., Mayes, M.A., Morrison, E., Riley, W.J., Salazar, A., Schimel, J.P., Tang, J., Classen, A.T., 2018. Multiple models and experiments underscore large uncertainty in soil carbon dynamics. Biogeochemistry 141, 109–123. doi:10.1007/s10533-018-0509-z

Sun, H., Santalahti, M., Pumpanen, J., Köster, K., Berninger, F., Raffaello, T., Jumpponen, A., Asiegbu, F.O., Heinonsalo, J., 2015. Fungal Community Shifts in Structure and Function across a Boreal Forest Fire Chronosequence. Applied and Environmental Microbiology 81, 7869–7880. doi:10.1128/AEM.02063-15

Szafrańska, K., Chowaniec, K., Dul, H., Zalewska-Gałosz, J., Skubała, K., 2026. Insight into the long-term impact of fire in dry pine forests on biological soil crust and underlying soil. Applied Soil Ecology 220, 106837. doi:10.1016/j.apsoil.2026.106837

Tao, F., Huang, Y., Hungate, B.A., Manzoni, S., Frey, S.D., Schmidt, M.W.I., Reichstein, M., Carvalhais, N., Ciais, P., Jiang, L., Lehmann, J., Wang, Y.-P., Houlton, B.Z., Ahrens, B., Mishra, U., Hugelius, G., Hocking, T.D., Lu, X., Shi, Z., Viatkin, K., Vargas, R., Yigini, Y., Omuto, C., Malik, A.A., Peralta, G., Cuevas-Corona, R., Di Paolo, L.E., Luotto, I., Liao, C., Liang, Y.-S., Saynes, V.S., Huang, X., Luo, Y., 2023. Microbial carbon use efficiency promotes global soil carbon storage. Nature 618, 981–985. doi:10.1038/s41586-023-06042-3

Tate, K.R., Ross, D.J., Feltham, C.W., 1988. A direct extraction method to estimate soil microbial c: effects of experimental variables and some different calibration procedures. Soil Biology and Biochemistry 20, 329–335. doi:10.1016/0038-0717(88)90013-2

Taylor, D.L., Walters, W.A., Lennon, N.J., Bochicchio, J., Krohn, A., Caporaso, J.G., Pennanen, T., 2016. Accurate estimation of fungal diversity and abundance through improved lineage-specific primers optimized for Illumina amplicon sequencing. Applied and Environmental Microbiology 82, 7217. doi:10.1128/AEM.02576-16

Ter-Mikaelian, M.T., Colombo, S.J., Chen, J., 2009. Estimating natural forest fire return interval in northeastern Ontario, Canada. Forest Ecology and Management 258, 2037–2045. doi:10.1016/j.foreco.2009.07.056

Thompson, D.K., Wotton, B.M., Waddington, J.M., 2015. Estimating the heat transfer to an organic soil surface during crown fire. International Journal of Wildland Fire 24, 120–129. doi:10.1071/WF12121

Treseder, K.K., Mack, M.C., Cross, A., 2004. Relationships among fires, fungi, and soil dynamics in Alaskan boreal forests. Ecological Applications 14, 1826–1838. doi:10.1890/03-5133

Úbeda, X., Bernia, S., Simelton, E., 2005. Chapter 6: The long-term effects on soil properties from a forest fire of varying intensity in a Mediterranean environment. Developments in Earth Surface Processes 7, 87–102. doi:10.1016/S0928-2025(05)80012-4

Waldrop, M.P., Harden, J.W., 2008. Interactive effects of wildfire and permafrost on microbial communities and soil processes in an Alaskan black spruce forest. Global Change Biology 14, 2591–2602. doi:10.1111/j.1365-2486.2008.01661.x

Walters, W., Hyde, E.R., Berg-Lyons, D., Ackermann, G., Humphrey, G., Parada, A., Gilbert, J.A., Jansson, J.K., Caporaso, J.G., Fuhrman, J.A., Apprill, A., Knight, R., 2015. Improved bacterial 16S rRNA gene (V4 and V4-5) and fungal internal transcribed spacer marker gene primers for microbial community surveys. mSystems 1. doi:10.1128/msystems.00009-15

Whitman, E., Parks, S.A., Holsinger, L.M., Parisien, M.A., 2022. Climate-induced fire regime amplification in Alberta, Canada. Environmental Research Letters 17, 055003. doi:10.1088/1748-9326/AC60D6

Whitman, T., Whitman, E., Woolet, J., Flannigan, M.D., Thompson, D.K., Parisien, M.-A., 2019. Soil bacterial and fungal response to wildfires in the Canadian boreal forest across a burn severity gradient. Soil Biology and Biochemistry 138, 107571. doi:10.1016/j.soilbio.2019.107571

Whitman, T., Woolet, J., Sikora, M., Johnson, D.B., Whitman, E., 2022. Resilience in soil bacterial communities of the boreal forest from one to five years after wildfire across a severity gradient. Soil Biology and Biochemistry 172, 108755. doi:10.1016/J.SOILBIO.2022.108755

Whitman, T., Woolet, J., Sikora, M.C., Johnson, D.B., Dawe, D.A., Whitman, E., 2025. Resilience not yet apparent in soil fungal communities of the boreal forest from one to five years after wildfire across a severity gradient. doi:10.1101/2025.03.29.646032

Wickham, H., 2016. ggplot2: Elegant graphics for data analysis. Springer-Verlag, New York.

Wickham, H., François, R., Henry, L., Müller, K., 2021. dplyr: A grammar of data manipulation. R package dplyr version 1.0.6.

Xiang, X., Shi, Y., Yang, J., Kong, J., Lin, X., Zhang, H., Zeng, J., Chu, H., 2014. Rapid recovery of soil bacterial communities after wildfire in a Chinese boreal forest. Scientific Reports 2014 4:1 4, 1–8. doi:10.1038/srep03829

Yano, K., Wada, T., Suzuki, S., Tagami, K., Matsumoto, T., Shiwa, Y., Ishige, T., Kawaguchi, Y., Masuda, K., Akanuma, G., Nanamiya, H., Niki, H., Yoshikawa, H., Kawamura, F., 2013. Multiple rRNA operons are essential for efficient cell growth and sporulation as well as outgrowth in *Bacillus subtilis*. Microbiology 159, 2225–2236. doi:10.1099/MIC.0.067025-0

Yilmaz, P., Parfrey, L.W., Yarza, P., Gerken, J., Pruesse, E., Quast, C., Schweer, T., Peplies, J., Ludwig, W., Glöckner, F.O., 2014. The SILVA and “All-species Living Tree Project (LTP)” taxonomic frameworks. Nucleic Acids Research 42, D643–D648. doi:10.1093/nar/gkt1209

Yin, J., Wu, D., Cooledge, E.C., Wang, C., Li, Y., Li, J., Wang, Q., Bol, R., Chadwick, D.R., Jones, D.L., 2026. Legacy Effects of Extreme Heat Decreased Soil Microbial Carbon Use Efficiency. Global Change Biology 32, e70967. doi:10.1111/gcb.70967

