## Supplemental information for "Persistent but variable effect of experimental laboratory burns on microbial community resistance, resilience, and function across contrasting boreal forest soils"

#### Methods: Pine-specific carbon use efficiency

After the 24-day incubation, the intact soil cores were destructively sampled. O and mineral horizons were separated, homogenized, and subsamples were collected for measurement of substrate-specific CUE using both $^{13}$C-labelled glucose and pine following the $^{13}$C-tracing method from Geyer et al. (2019). Four subsamples were collected from each soil horizon and placed in 125 mL Mason jars. $^{13}$C-labelled glucose (5 at%) or $^{13}$C-labelled pine (1.6 at%) was then added to two of the jars and unlabeled glucose (1.1 at%) or pine (1.1 at%) was added to the remaining two jars at a rate of 1.0 mg glucose $g^{-1}$ dry soil (0.4 mg C $g^{-1}$ dry soil) or 2.4 mg pine $g^{-1}$ dry soil (0.91 mg C $g^{-1}$ dry soil), respectively (Table S3), and soil was mixed well. The pine amendment was created by finely grinding pulse-labelled roots of eastern white pine (*Pinus strobus* (L)). Briefly, we used the belowground biomass from two-year-old *Pinus strobus* tree seedlings grown in two custom growth chambers for one growing season under controlled moisture, humidity, temperature, and light conditions. In the first chamber, trees were pulse labeled with 99% $^{13}CO_2$ at regular intervals ("labelled pine"). In the second chamber, trees were exposed to ambient levels of non-enriched $CO_2$ ("unlabeled pine") (full details in Zeba et al. 2024).

#### Carbon use efficiency calculations and derivatizations

Adjust dissolved organic C (DOC) concentrations following Equations S1 & S2:

$$DOC_{F,adj.} = DOC_F - DOC_{F,blank} \quad \text{Eq. S1}$$

$$DOC_{NF,adj.} = DOC_{NF} - DOC_{NF,blank} \quad \text{Eq. S2}$$

where $DOC_F$ and $DOC_{NF}$ represent the total dissolved organic carbon (µg C $g^{-1}$ dry soil) from fumigated (F) and non-fumigated (NF) soils, respectively, and $DOC_{F,adj.}$ and $DOC_{NF,adj.}$ represent blank-adjusted total dissolved organic carbon.

Calculate total microbial biomass C using Equation S3:

$$MBC = DOC_{F,\,adj} - DOC_{NF,adj} \quad \text{Eq. S3}$$

where MBC represents total microbial biomass C (μg C $g^{-1}$ dry soil).

Calculating the atom % of $^{13}$C in MBC from soils with labelled and unlabeled amendments using Equations S4 and S5, respectively:

$$\text{at\% MBC}_{soil+LA} = \frac{(\text{at\%DOC}_{F,LA} \times [\text{DOC}_{F,LA}]) - (\text{at\%DOC}_{NF,LA} \times [\text{DOC}_{NF,LA}])}{(\text{DOC}_{F,LA} - \text{DOC}_{NF,LA})} \qquad \text{Eq. S4}$$

$$\text{at\% MBC}_{soil+UA} = \frac{(\text{at\%DOC}_{F,UA} \times [\text{DOC}_{F,UA}]) - (\text{at\%DOC}_{NF,UA} \times [\text{DOC}_{NF,UA}])}{(\text{DOC}_{F,UA} - \text{DOC}_{NF,UA})} \qquad \text{Eq. S5}$$

where at% $\text{MBC}_{soil+LA}$ represent the atom % of $^{13}$C in total MBC from soils with the labelled amendment (LA), and at% $\text{DOC}_{F,LA}$ and at% $\text{DOC}_{NF,LA}$ represent the atom % of $^{13}$C in DOC from soils with the labelled amendment that were fumigated or non-fumigated, respectively. $\text{DOC}_{F,LA}$ and $\text{DOC}_{NF,LA}$ represent the DOC (μg C $g^{-1}$ dry soil) from fumigated and non-fumigated soils that were first amended with the labelled substrate. At% MBCsoil+$_{UA}$ represents the atom % of $^{13}$C in total MBC from soils with the unlabeled amendment (UA), and at% $\text{DOC}_{F,UA}$ and at% $\text{DOC}_{NF,UA}$ represent the atom % of $^{13}$C in DOC from soils with the unlabeled amendment that were fumigated or non-fumigated, respectively. $\text{DOC}_{F,UA}$ and $\text{DOC}_{NF,UA}$ represent the DOC (μg C $g^{-1}$ dry soil) from fumigated and non-fumigated soils that were first amended with the unlabeled substrate.

Calculate the fraction of total MBC that is substrate-derived using Equation S6:

$$f_{MBC,LA} = \frac{(\text{at\%MBC}_{soil+LA} - \text{at\%MBC}_{soil+UA})}{(\text{at\%LA} - \text{at\%UA})} \qquad \text{Eq. S6}$$

where $f_{MBC,LA}$ is the fraction of total MBC derived from the labelled amendment, and at% LA and at% UA is the atom % of $^{13}$C in the labelled and unlabeled amendments, respectively.

<u>Derivation of Equation S6</u>

Starting equations:

$$\text{at\%MBC}_{soil+LA} = f_{MBC,LA} \times \text{at\%LA} + f_{MBC,soil} \times \text{at\%soil}$$
$$\text{at\%MBC}_{soil+UA} = f_{MBC,UA} \times \text{at\%UA} + f_{MBC,soil} \times \text{at\%soil}$$

Assumption: $f_{MBC,LA} \approx f_{MBC,UA}$

Derivation steps

1. $f_{MBC,soil} \times \text{at\%soil} = \text{at\%MBC}_{soil+LA} - f_{MBC,LA} \times \text{at\%LA}$
2. $f_{MBC,soil} \times \text{at\%soil} = \text{at\%MBC}_{soil+UA} - f_{MBC,UA} \times \text{at\%UA}$
3. $f_{MBC,soil} \times \text{at\%soil} = \text{at\%MBC}_{soil+UA} - f_{MBC,LA} \times \text{at\%UA}$
4. $\text{at\%MBC}_{soil+LA} - f_{MBC,LA} \times \text{at\%LA} = \text{at\%MBC}_{soil+UA} - f_{MBC,LA} \times \text{at\%UA}$
5. $\text{at\%MBC}_{soil+LA} - \text{at\%MBC}_{soil+UA} = f_{MBC,LA} \times \text{at\%LA} - f_{MBC,LA} \times \text{at\%UA}$
6. $\text{at\%MBC}_{soil+LA} - \text{at\%MBC}_{soil+UA} = f_{MBC,LA} \times (\text{at\%LA} - \text{at\%UA})$
7. $f_{MBC,LA} = \frac{(\text{at\%MBC}_{soil+LA} - \text{at\%MBC}_{soil+UA})}{(\text{at\%LA} - \text{at\%UA})}$

Calculating the total substrate-derived MBC (μg C $g^{-1}$ dry soil) using Equation S7:

$$\mathrm{MBC_{LA}} = \mathrm{MBC} \times \mathrm{f_{MBC,LA}} \qquad \text{Eq. S7}$$

where $MBC_{LA}$ represents substrate-derived MBC (μg C $g^{-1}$ dry soil).

Calculating the fraction of total $CO_2$ that is derived from the labelled substrate using Equation S8:

$$\mathrm{f_{CO2,LA}} = \frac{\delta^{13}\mathrm{C_{CO2,soil+LA}} - \delta^{13}\mathrm{C_{CO2,soil+UA}}}{\delta^{13}\mathrm{C_{LA}} - \delta^{13}\mathrm{C_{UA}}} \qquad \text{Eq. S8}$$

where $f_{CO2,LA}$ is the fraction of total $CO_2$ derived from labelled amendment, $\delta^{13}C_{CO2,soil+LA}$ and $\delta^{13}C_{CO2,soil+UA}$ represent the $\delta^{13}C$ values of $CO_2$ from soils and the labelled or unlabeled amendment, respectively, and $\delta^{13}C_{LA}$ and $\delta^{13}C_{UA}$ represent the $\delta^{13}C$ values of the labelled and unlabeled amendments, respectively.

<u>Derivation of Equation S8</u>

Starting equations:

$$\delta^{13}C_{CO2,soil+LA} = f_{CO2,soil} \times \delta^{13}C_{soil} + f_{CO2,LA} \times \delta^{13}C_{LA}$$
$$\delta^{13}C_{CO2,soil+UA} = f_{CO2,soil} \times \delta^{13}C_{soil} + f_{CO2,UA} \times \delta^{13}C_{UA}$$

Assumption: $f_{LA} = f_{UA}$

Where $\delta^{13}C_{CO2,soil+LA}$ and $\delta^{13}C_{CO2,soil+UA}$ represent the $\delta^{13}C$ of $CO_2$ from soils with the labelled or unlabeled amendment, respectively; $f_{CO2,soil}$ and $\delta^{13}C_{soil}$ represent the fraction of total $CO_2$-C derived from soil and the $\delta^{13}C$ signature of the soil, respectively; $f_{CO2,LA}$ and $f_{CO2,UA}$ represent the fraction of total $CO_2$-C derived from the labelled and unlabeled amendments, respectively; and $\delta^{13}C_{LA}$ and $\delta^{13}C_{UA}$ represent the $\delta^{13}C$ signature of the labelled and unlabeled amendment, respectively.

Calculate total substrate-derived $CO_2$ (μg C $g^{-1}$ dry soil) using Equation S9:

$$\mathrm{CO_{2\,LA}} = \mathrm{CO_{2\,total}} \times \mathrm{f_{CO2,LA}} \qquad \text{Eq. S9}$$

where $CO_{2\,LA}$ represents substrate-derived $CO_2$ (μg C $g^{-1}$ dry soil), and $CO_{2\,total}$ represents total $CO_2$ respired (μg C $g^{-1}$ dry soil).

Calculate CUE using Equation S10:

$$\mathrm{CUE} = \frac{\mathrm{MBC_{LA}}}{(\mathrm{MBC_{LA}} + \mathrm{CO_{2,LA}})} = \frac{\text{substrate derived MBC}}{(\text{substrate derived MBC} + \text{substrate derived } \mathrm{CO_2})} \qquad \text{Eq. S10}$$

Calculate the metabolic quotient, $qCO_2$, using Equation S11:

$$\mathrm{qCO_2} = \frac{\mathrm{CO_{2,LA}}}{\mathrm{MBC_{LA}}} \qquad \text{Eq. S11}$$

### Supplementary Tables

Table S1. Site locations and characteristics.

| Site ID | Latitude | Longitude | Soil texture class | Soil type | Dominant tree species |
|---|---|---|---|---|---|
| 1 | -112.26 | 59.49 | Loamy Sand | Gleysol | *Pinus banksiana* |
| 2 | -112.41 | 59.40 | Sand | Gleysol | *Pinus banksiana* |
| 3 | -112.49 | 59.33 | Sand | Gleysol | *Pinus banksiana* |
| 5 | -112.39 | 59.41 | Sandy Loam | Gleysol | *Pinus banksiana* |
| 8 | -112.48 | 59.36 | Silt Loam | Gleysol | *Pinus banksiana* |
| 9 | -112.41 | 59.38 | Sand | Gleysol | *Pinus banksiana* |
| 4 | -112.49 | 59.31 | Organic | Histosol | *Picea* spp. |
| 6 | -112.25 | 59.51 | Organic | Histosol | *Picea* spp. |
| 7 | -112.36 | 59.44 | Organic | Histosol | *Picea* spp. |
| 10 | -112.42 | 59.39 | Organic | Histosol | *Picea* spp. |
| 11 | -112.38 | 59.43 | Organic | Histosol | *Picea* spp. |
| 12 | -112.49 | 59.33 | Organic | Histosol | *Picea* spp. |

Table S2. Full 16S PCR primers with barcodes.

| Primer Name | Sequence |
|---|---|
| 515f_SA501 | AATGATACGGCGACCACCGAGATCTACACATCGTACGTATGGTAATTGTGTGYCAGCMGCCGCGGTAA |
| 515f_SA502 | AATGATACGGCGACCACCGAGATCTACACACTATCTGTATGGTAATTGTGTGYCAGCMGCCGCGGTAA |
| 515f_SA503 | AATGATACGGCGACCACCGAGATCTACACTAGCGAGTTATGGTAATTGTGTGYCAGCMGCCGCGGTAA |
| 515f_SA504 | AATGATACGGCGACCACCGAGATCTACACCTGCGTGTTATGGTAATTGTGTGYCAGCMGCCGCGGTAA |
| 515f_SA505 | AATGATACGGCGACCACCGAGATCTACACTCATCGAGTATGGTAATTGTGTGYCAGCMGCCGCGGTAA |
| 515f_SA506 | AATGATACGGCGACCACCGAGATCTACACCGTGAGTGTATGGTAATTGTGTGYCAGCMGCCGCGGTAA |
| 515f_SA507 | AATGATACGGCGACCACCGAGATCTACACGGATATCTTATGGTAATTGTGTGYCAGCMGCCGCGGTAA |
| 515f_SA508 | AATGATACGGCGACCACCGAGATCTACACGACACCGTTATGGTAATTGTGTGYCAGCMGCCGCGGTAA |
| 515f_SB501 | AATGATACGGCGACCACCGAGATCTACACCTACTATATATGGTAATTGTGTGYCAGCMGCCGCGGTAA |
| 515f_SB502 | AATGATACGGCGACCACCGAGATCTACACCGTTACTATATGGTAATTGTGTGYCAGCMGCCGCGGTAA |
| 515f_SB503 | AATGATACGGCGACCACCGAGATCTACACAGAGTCACTATGGTAATTGTGTGYCAGCMGCCGCGGTAA |
| 515f_SB504 | AATGATACGGCGACCACCGAGATCTACACTACGAGACTATGGTAATTGTGTGYCAGCMGCCGCGGTAA |
| 515f_SB505 | AATGATACGGCGACCACCGAGATCTACACACGTCTCGTATGGTAATTGTGTGYCAGCMGCCGCGGTAA |
| 515f_SB506 | AATGATACGGCGACCACCGAGATCTACACTCGACGAGTATGGTAATTGTGTGYCAGCMGCCGCGGTAA |
| 515f_SB507 | AATGATACGGCGACCACCGAGATCTACACGATCGTGTTATGGTAATTGTGTGYCAGCMGCCGCGGTAA |
| 515f_SB508 | AATGATACGGCGACCACCGAGATCTACACGTCAGATATATGGTAATTGTGTGYCAGCMGCCGCGGTAA |
| 515f_SC501 | AATGATACGGCGACCACCGAGATCTACACACGACGTGTATGGTAATTGTGTGYCAGCMGCCGCGGTAA |
| 515f_SC502 | AATGATACGGCGACCACCGAGATCTACACATATACACTATGGTAATTGTGTGYCAGCMGCCGCGGTAA |
| 515f_SC503 | AATGATACGGCGACCACCGAGATCTACACCGTCGCTATATGGTAATTGTGTGYCAGCMGCCGCGGTAA |

| | |
|---|---|
| 515f_SC504 | AATGATACGGCGACCACCGAGATCTACACCTAGAGCTTATGGTAATTGTGTGYCAGCMGCCGCGGTAA |
| 515f_SC505 | AATGATACGGCGACCACCGAGATCTACACGCTCTAGTTATGGTAATTGTGTGYCAGCMGCCGCGGTAA |
| 515f_SC506 | AATGATACGGCGACCACCGAGATCTACACGACACTGATATGGTAATTGTGTGYCAGCMGCCGCGGTAA |
| 515f_SC507 | AATGATACGGCGACCACCGAGATCTACACTGCGTACGTATGGTAATTGTGTGYCAGCMGCCGCGGTAA |
| 515f_SC508 | AATGATACGGCGACCACCGAGATCTACACTAGTGTAGTATGGTAATTGTGTGYCAGCMGCCGCGGTAA |
| 806r_SA701 | CAAGCAGAAGACGGCATACGAGATAACTCTCGAGTCAGCCAGCCGGACTACNVGGGTWTCTAAT |
| 806r_SA702 | CAAGCAGAAGACGGCATACGAGATACTATGTCAGTCAGCCAGCCGGACTACNVGGGTWTCTAAT |
| 806r_SA703 | CAAGCAGAAGACGGCATACGAGATAGTAGCGTAGTCAGCCAGCCGGACTACNVGGGTWTCTAAT |
| 806r_SA704 | CAAGCAGAAGACGGCATACGAGATCAGTGAGTAGTCAGCCAGCCGGACTACNVGGGTWTCTAAT |
| 806r_SA705 | CAAGCAGAAGACGGCATACGAGATCGTACTCAAGTCAGCCAGCCGGACTACNVGGGTWTCTAAT |
| 806r_SA706 | CAAGCAGAAGACGGCATACGAGATCTACGCAGAGTCAGCCAGCCGGACTACNVGGGTWTCTAAT |
| 806r_SA707 | CAAGCAGAAGACGGCATACGAGATGGAGACTAAGTCAGCCAGCCGGACTACNVGGGTWTCTAAT |
| 806r_SA708 | CAAGCAGAAGACGGCATACGAGATGTCGCTCGAGTCAGCCAGCCGGACTACNVGGGTWTCTAAT |
| 806r_SA709 | CAAGCAGAAGACGGCATACGAGATGTCGTAGTAGTCAGCCAGCCGGACTACNVGGGTWTCTAAT |
| 806r_SA710 | CAAGCAGAAGACGGCATACGAGATTAGCAGACAGTCAGCCAGCCGGACTACNVGGGTWTCTAAT |
| 806r_SA711 | CAAGCAGAAGACGGCATACGAGATTCATAGACAGTCAGCCAGCCGGACTACNVGGGTWTCTAAT |
| 806r_SA712 | CAAGCAGAAGACGGCATACGAGATTCGCTATAAGTCAGCCAGCCGGACTACNVGGGTWTCTAAT |
| 806r_SB701 | CAAGCAGAAGACGGCATACGAGATAAGTCGAGAGTCAGCCAGCCGGACTACNVGGGTWTCTAAT |
| 806r_SB702 | CAAGCAGAAGACGGCATACGAGATATACTTCGAGTCAGCCAGCCGGACTACNVGGGTWTCTAAT |
| 806r_SB703 | CAAGCAGAAGACGGCATACGAGATAGCTGCTAAGTCAGCCAGCCGGACTACNVGGGTWTCTAAT |
| 806r_SB704 | CAAGCAGAAGACGGCATACGAGATCATAGAGAAGTCAGCCAGCCGGACTACNVGGGTWTCTAAT |
| 806r_SB705 | CAAGCAGAAGACGGCATACGAGATCGTAGATCAGTCAGCCAGCCGGACTACNVGGGTWTCTAAT |
| 806r_SB706 | CAAGCAGAAGACGGCATACGAGATCTCGTTACAGTCAGCCAGCCGGACTACNVGGGTWTCTAAT |
| 806r_SB707 | CAAGCAGAAGACGGCATACGAGATGCGCACGTAGTCAGCCAGCCGGACTACNVGGGTWTCTAAT |
| 806r_SB708 | CAAGCAGAAGACGGCATACGAGATGGTACTATAGTCAGCCAGCCGGACTACNVGGGTWTCTAAT |
| 806r_SB709 | CAAGCAGAAGACGGCATACGAGATGTATACGCAGTCAGCCAGCCGGACTACNVGGGTWTCTAAT |
| 806r_SB710 | CAAGCAGAAGACGGCATACGAGATTACGAGCAAGTCAGCCAGCCGGACTACNVGGGTWTCTAAT |
| 806r_SB711 | CAAGCAGAAGACGGCATACGAGATTCAGCGTTAGTCAGCCAGCCGGACTACNVGGGTWTCTAAT |
| 806r_SB712 | CAAGCAGAAGACGGCATACGAGATTCGCTACGAGTCAGCCAGCCGGACTACNVGGGTWTCTAAT |

Table S3. Full ITS2 PCR primers with barcodes.

| Primer Name | Primer Sequence |
|---|---|
| ITS4-6 | AATGATACGGCGACCACCGAGATCTACACCCAAGCTATATGGTAATTAAAGCCTCCGCTTATTGATATGCTTAART |
| ITS4-7 | AATGATACGGCGACCACCGAGATCTACACGGCTCGATTATGGTAATTAAAGCCTCCGCTTATTGATATGCTTAART |
| ITS4-13 | AATGATACGGCGACCACCGAGATCTACACCGTAGACTTATGGTAATTAAAGCCTCCGCTTATTGATATGCTTAART |
| ITS4-15 | AATGATACGGCGACCACCGAGATCTACACGGATAAGGTATGGTAATTAAAGCCTCCGCTTATTGATATGCTTAART |
| ITS4-17 | AATGATACGGCGACCACCGAGATCTACACTTCTAAGTTATGGTAATTAAAGCCTCCGCTTATTGATATGCTTAART |
| ITS4-21 | AATGATACGGCGACCACCGAGATCTACACACGCATTCTATGGTAATTAAAGCCTCCGCTTATTGATATGCTTAART |
| ITS4-25 | AATGATACGGCGACCACCGAGATCTACACTGCCTACTTATGGTAATTAAAGCCTCCGCTTATTGATATGCTTAART |
| ITS4-27 | AATGATACGGCGACCACCGAGATCTACACATTATGGTTATGGTAATTAAAGCCTCCGCTTATTGATATGCTTAART |
| ITS4-28 | AATGATACGGCGACCACCGAGATCTACACCTTAGCAGTATGGTAATTAAAGCCTCCGCTTATTGATATGCTTAART |
| ITS4-34 | AATGATACGGCGACCACCGAGATCTACACTCTTGAATTATGGTAATTAAAGCCTCCGCTTATTGATATGCTTAART |
| 5.8S-A | CAAGCAGAAGACGGCATACGAGATCATGAGAGAGTCAGTCAGGGAACTTTYRRCAAYGGATCWCT |
| 5.8S-B | CAAGCAGAAGACGGCATACGAGATTTGGCAACAGTCAGTCAGGGAACTTTYRRCAAYGGATCWCT |
| 5.8S-E | CAAGCAGAAGACGGCATACGAGATTTCCAGCAAGTCAGTCAGGGAACTTTYRRCAAYGGATCWCT |
| 5.8S-G | CAAGCAGAAGACGGCATACGAGATCCAATTGCAGTCAGTCAGGGAACTTTYRRCAAYGGATCWCT |
| 5.8S-H | CAAGCAGAAGACGGCATACGAGATCTCATGCGAGTCAGTCAGGGAACTTTYRRCAAYGGATCWCT |
| 5.8S-J | CAAGCAGAAGACGGCATACGAGATCTGCAGACAGTCAGTCAGGGAACTTTYRRCAAYGGATCWCT |
| 5.8S-K | CAAGCAGAAGACGGCATACGAGATTGCTATAAAGTCAGTCAGGGAACTTTYRRCAAYGGATCWCT |
| 5.8S-L | CAAGCAGAAGACGGCATACGAGATAACGTTGGAGTCAGTCAGGGAACTTTYRRCAAYGGATCWCT |
| 5.8S-M | CAAGCAGAAGACGGCATACGAGATCCGGAACTAGTCAGTCAGGGAACTTTYRRCAAYGGATCWCT |
| 5.8S-N | CAAGCAGAAGACGGCATACGAGATCTTCAACGAGTCAGTCAGGGAACTTTYRRCAAYGGATCWCT |
| 5.8S-Q | CAAGCAGAAGACGGCATACGAGATGCATTCCAAGTCAGTCAGGGAACTTTYRRCAAYGGATCWCT |
| 5.8S-R | CAAGCAGAAGACGGCATACGAGATCGCTGGCAAGTCAGTCAGGGAACTTTYRRCAAYGGATCWCT |
| 5.8S-T | CAAGCAGAAGACGGCATACGAGATCGCGCAGTAGTCAGTCAGGGAACTTTYRRCAAYGGATCWCT |
| 5.8S-W | CAAGCAGAAGACGGCATACGAGATTTCTGGACAGTCAGTCAGGGAACTTTYRRCAAYGGATCWCT |
| 5.8S-X | CAAGCAGAAGACGGCATACGAGATTTCAGCCTAGTCAGTCAGGGAACTTTYRRCAAYGGATCWCT |
| 5.8S-Y | CAAGCAGAAGACGGCATACGAGATCTATATTGAGTCAGTCAGGGAACTTTYRRCAAYGGATCWCT |
| 5.8S-Z | CAAGCAGAAGACGGCATACGAGATCTCGTTAAAGTCAGTCAGGGAACTTTYRRCAAYGGATCWCT |

Table S4. Mass of amendment C added to soil for incubation and mass of chloroform used during microbial biomass C measurement.

| Addition to soil | Soil type | C added (mg per g dry soil) | C added (percent of total C) |
|---|---|---|---|
| Glucose | Histosol | 0.4 ± 0.002 | 0.11% |
| | O horizon, Gleysol | 0.4 ± 0.005 | 0.23% |
| | Mineral soil, Gleysol | 0.4 ± 0.0008 | 2.9% |
| Pine | Histosol | 1.0 ± 0.005 | 0.27% |
| | O horizon, Gleysol | 1.0 ± 0.01 | 0.57% |
| | Mineral soil, Gleysol | 1.0 ± 0.002 | 7.2% |
| Chloroform* | Histosol | 109 ± 2 | 29% |
| | O horizon, Gleysol | 120 ± 55 | 75% |
| | Mineral soil, Gleysol | 17 ± 0.4 | 120% |

* Traces of chloroform were removed from the extracted solution by bubbling with lab air before analysis

Table S5. PERMANOVA results for full model from 16S rRNA gene.

| | Df | Sum of Squares | $R^2$ | F | Pr(>F) |
|---|---|---|---|---|---|
| Burn treatment duration | 2 | 2.18 | 0.0302 | 4.91 | 0.001 |
| Soil type and horizon | 2 | 20.1 | 0.279 | 45.3 | 0.001 |
| pH | 1 | 1.90 | 0.0263 | 8.55 | 0.001 |
| Incubation length | 3 | 1.86 | 0.0257 | 2.79 | 0.001 |
| Degree hours (at uppermost thermocouple) | 1 | 1.48 | 0.0206 | 6.68 | 0.001 |
| Residual | 201 | 44.6 | 0.619 | | |
| Total | 210 | 72.1 | 1.00 | | |

Table S6. Single-component PERMANOVA model results for 16S rRNA gene. The order of terms in the PERMANOVA model affects the partial $R^2$ for a given term; thus, to compare the relative explanatory power of each component, we report the $R^2$ of single-component models for each factor included into the full 16S PERMANOVA model (Table S5).

| Factor | $R^2$ | Pr(>F) |
|---|---|---|
| Burn treatment duration | 0.030 | 0.001 |
| Soil type and horizon | 0.28 | 0.001 |
| pH | 0.17 | 0.001 |
| Incubation length | 0.023 | 0.011 |
| Degree hours | 0.050 | 0.001 |

Table S7. PERMNANOVA results for full model from ITS gDNA.

| | Df | Sum of Squares | $R^2$ | F | Pr(>F) |
|---|---|---|---|---|---|
| Burn treatment duration | 2 | 3.60 | 0.0409 | 5.24 | 0.001 |
| Soil type and horizon | 2 | 12.2 | 0.138 | 17.7 | 0.001 |
| pH | 1 | 1.07 | 0.0121 | 3.11 | 0.001 |
| Incubation length | 3 | 2.29 | 0.0259 | 2.21 | 0.001 |
| Degree hours (at uppermost thermocouple) | 1 | 1.32 | 0.0150 | 3.84 | 0.001 |
| Residual | 197 | 67.8 | 0.768 | | |
| Total | 206 | 88.2 | 1.00 | | |

Table S8. Single-component PERMANOVA model results for ITS gDNA gene. The order of terms in the PERMANOVA model affects the partial $R^2$ for a given term; thus, to compare the relative explanatory power of each component, we report the $R^2$ of single-component models for each factor included into the full ITS PERMANOVA model (Table S7).

| Factor | $R^2$ | Pr(>F) |
|---|---|---|
| Burn treatment duration | 0.041 | 0.001 |
| Soil type and horizon | 0.14 | 0.001 |
| pH | 0.056 | 0.001 |
| Incubation length | 0.026 | 0.001 |
| Degree hours | 0.030 | 0.001 |

Table S9. Weighted mean predicted 16S rRNA gene copy number for all samples.

| Sample ID | **Weighted mean predicted 16S rRNA gene copy number** | |
|---|---|---|
| | O horizon | Mineral soil |
| 22UW-WB-01-03 | 4.57 | 1.93 |
| 22UW-WB-01-04 | 2.17 | 1.74 |
| 22UW-WB-01-05 | 1.65 | 1.59 |
| 22UW-WB-01-06 | 1.87 | 1.64 |
| 22UW-WB-01-07 | 1.85 | 1.64 |
| 22UW-WB-01-09 | 1.81 | 1.50 |
| 22UW-WB-01-11 | 1.87 | 1.62 |
| 22UW-WB-01-12 | 3.35 | 1.73 |
| 22UW-WB-01-13 | 3.11 | 1.88 |
| 22UW-WB-01-15 | 1.83 | 1.67 |
| 22UW-WB-01-17 | 3.72 | 1.73 |
| 22UW-WB-01-18 | 1.56 | 1.67 |
| 22UW-WB-02-02 | 4.59 | 1.79 |
| 22UW-WB-02-03 | 3.30 | 1.75 |
| 22UW-WB-02-04 | 2.88 | 1.70 |
| 22UW-WB-02-05 | 1.98 | 2.19 |
| 22UW-WB-02-06 | 1.97 | 1.60 |
| 22UW-WB-02-09 | 3.05 | 1.80 |
| 22UW-WB-02-11 | 2.97 | 1.74 |
| 22UW-WB-02-12 | 2.07 | 1.55 |
| 22UW-WB-02-13 | 2.45 | 1.66 |
| 22UW-WB-02-15 | 9.48 | 1.93 |
| 22UW-WB-02-16 | 1.73 | 1.64 |
| 22UW-WB-02-18 | 2.14 | 1.69 |
| 22UW-WB-03-02 | 1.91 | 1.64 |
| 22UW-WB-03-04 | 2.15 | 1.61 |
| 22UW-WB-03-05 | 1.58 | 1.52 |
| 22UW-WB-03-06 | 1.71 | 1.51 |
| 22UW-WB-03-07 | 2.02 | |
| 22UW-WB-03-08 | 2.02 | 1.73 |
| 22UW-WB-03-09 | 3.38 | 1.86 |
| 22UW-WB-03-11 | 2.22 | 1.95 |
| 22UW-WB-03-12 | 2.10 | 1.70 |
| 22UW-WB-03-15 | 1.68 | 1.55 |
| 22UW-WB-03-16 | 1.80 | 1.60 |
| 22UW-WB-03-17 | 2.57 | 1.57 |
| 22UW-WB-04-02 | 1.64 | |
| 22UW-WB-04-03 | 1.57 | |
| 22UW-WB-04-04 | 1.95 | |
| 22UW-WB-04-05 | 1.59 | |
| 22UW-WB-04-06 | 1.87 | |
| 22UW-WB-04-07 | 1.93 | |
| 22UW-WB-04-08 | 1.70 | |
| 22UW-WB-04-09 | 1.57 | |
| 22UW-WB-04-13 | 1.73 | |
| 22UW-WB-04-15 | 1.67 | |
| 22UW-WB-04-16 | 1.53 | |
| 22UW-WB-04-17 | 1.63 | |
| 22UW-WB-05-02 | 1.68 | 1.63 |
| 22UW-WB-05-05 | 1.71 | 1.57 |
| 22UW-WB-05-06 | 1.62 | 1.63 |
| 22UW-WB-05-07 | 2.07 | 1.71 |
| 22UW-WB-05-08 | 3.68 | 2.41 |
| 22UW-WB-05-09 | 1.85 | 1.50 |
| 22UW-WB-05-11 | 2.62 | 1.75 |
| 22UW-WB-05-12 | 1.74 | 1.55 |
| 22UW-WB-05-15 | | 1.65 |
| 22UW-WB-05-16 | | 1.53 |
| 22UW-WB-05-17 | 1.74 | 1.63 |
| 22UW-WB-05-18 | 1.99 | 1.60 |
| 22UW-WB-06-02 | 2.67 | |
| 22UW-WB-06-03 | 1.56 | |
| 22UW-WB-06-04 | 1.71 | |
| 22UW-WB-06-05 | 1.60 | |
| 22UW-WB-06-06 | 1.62 | |
| 22UW-WB-06-07 | 2.01 | |
| 22UW-WB-06-08 | 1.72 | |
| 22UW-WB-06-11 | 2.20 | |
| 22UW-WB-06-12 | 1.62 | |
| 22UW-WB-06-13 | 2.30 | |
| 22UW-WB-06-15 | 1.68 | |
| 22UW-WB-06-17 | 1.76 | |
| 22UW-WB-07-04 | 2.23 | |
| 22UW-WB-07-05 | 2.30 | |
| 22UW-WB-07-06 | 2.20 | |
| 22UW-WB-07-07 | 1.55 | |
| 22UW-WB-07-08 | 1.71 | |
| 22UW-WB-07-11 | 2.35 | |
| 22UW-WB-07-12 | 2.33 | |

| | | |
|---|---|---|
| 22UW-WB-07-13 | 2.41 | |
| 22UW-WB-07-15 | 1.88 | |
| 22UW-WB-07-16 | 1.73 | |
| 22UW-WB-07-17 | 1.74 | |
| 22UW-WB-07-18 | 1.96 | |
| 22UW-WB-08-02 | 2.47 | 1.66 |
| 22UW-WB-08-04 | 1.83 | 2.08 |
| 22UW-WB-08-06 | 2.14 | |
| 22UW-WB-08-07 | 3.42 | 1.80 |
| 22UW-WB-08-08 | 1.88 | 1.62 |
| 22UW-WB-08-09 | 2.04 | 1.67 |
| 22UW-WB-08-11 | 2.08 | 1.63 |
| 22UW-WB-08-12 | 2.25 | 1.73 |
| 22UW-WB-08-13 | 2.26 | 1.69 |
| 22UW-WB-08-15 | 2.69 | 1.68 |
| 22UW-WB-08-16 | 4.85 | 2.85 |
| 22UW-WB-08-18 | 1.82 | 1.68 |
| 22UW-WB-09-02 | 1.97 | 1.57 |
| 22UW-WB-09-03 | 1.97 | 1.59 |
| 22UW-WB-09-04 | 1.48 | 1.64 |
| 22UW-WB-09-05 | 1.73 | 1.60 |
| 22UW-WB-09-06 | 1.64 | 1.75 |
| 22UW-WB-09-07 | 3.41 | 1.81 |
| 22UW-WB-09-12 | 2.99 | 1.71 |
| 22UW-WB-09-13 | 1.61 | 1.54 |
| 22UW-WB-09-15 | 1.64 | 1.62 |
| 22UW-WB-09-16 | 1.93 | 1.60 |
| 22UW-WB-09-17 | 1.62 | 1.59 |
| 22UW-WB-09-18 | 1.73 | 1.66 |
| 22UW-WB-11-02 | 1.54 | |
| 22UW-WB-11-03 | 1.98 | |
| 22UW-WB-11-04 | 1.76 | |
| 22UW-WB-11-06 | 2.47 | |
| 22UW-WB-11-07 | 3.37 | |
| 22UW-WB-11-08 | 1.58 | |
| 22UW-WB-11-09 | 1.55 | |
| 22UW-WB-11-12 | 2.10 | |
| 22UW-WB-11-15 | 1.57 | |
| 22UW-WB-11-16 | 1.82 | |
| 22UW-WB-11-17 | 2.74 | |
| 22UW-WB-11-18 | 1.55 | |
| 22UW-WB-12-02 | 1.78 | |
| 22UW-WB-12-03 | 2.15 | |
| 22UW-WB-12-04 | 1.64 | |
| 22UW-WB-12-05 | 1.70 | |
| 22UW-WB-12-07 | 1.76 | |
| 22UW-WB-12-08 | 3.37 | |
| 22UW-WB-12-09 | 1.82 | |
| 22UW-WB-12-11 | 1.62 | |
| 22UW-WB-12-12 | 1.75 | |
| 22UW-WB-12-13 | 2.45 | |
| 22UW-WB-12-15 | 1.58 | |
| 22UW-WB-12-18 | 1.72 | |

Table S10. Weighted mean predicted 16S rRNA gene copy number averaged across soil type, horizon, burn treatment duration, and incubation length.

| **Soil type** | **Horizon** | **Burn treatment duration (s)** | **Incubation length (days)** | **Predicted weighted mean 16S rRNA gene copy number (mean ± sd)** |
|---|---|---|---|---|
| Histosol | O | 0 | 2 | 1.63 ± 0.074 |
| | | 0 | 24 | 1.63 ± 0.072 |
| | | 0 | 49 | 1.67 ± 0.12 |
| | | 0 | 70 | 1.63 ± 0.13 |
| | | 30 | 2 | 2.04 ± 0.48 |
| | | 30 | 24 | 1.77 ± 0.32 |
| | | 30 | 49 | 1.77 ± 0.17 |
| | | 30 | 70 | 1.75 ± 0.23 |
| | | 120 | 2 | 2.31 ± 0.60 |
| | | 120 | 24 | 2.57 ± 0.73 |
| | | 120 | 49 | 2.27 ± 0.70 |
| | | 120 | 70 | 2.01 ± 0.29 |
| Gleysol | O | 0 | 2 | 1.71 ± 0.13 |
| | | 0 | 24 | 1.85 ± 0.15 |
| | | 0 | 49 | 1.75 ± 0.18 |
| | | 0 | 70 | 1.88 ± 0.14 |
| | | 30 | 2 | 2.27 ± 0.69 |
| | | 30 | 24 | 2.71 ± 1.12 |
| | | 30 | 49 | 2.01 ± 0.38 |
| | | 30 | 70 | 1.92 ± 0.22 |
| | | 120 | 2 | 3.85 ± 2.97 |
| | | 120 | 24 | 3.37 ± 0.80 |
| | | 120 | 49 | 3.15 ± 0.41 |
| | | 120 | 70 | 2.21 ± 0.31 |
| | M | 0 | 2 | 1.68 ± 0.20 |
| | | 0 | 24 | 1.62 ± 0.049 |
| | | 0 | 49 | 1.61 ± 0.060 |
| | | 0 | 70 | 1.69 ± 0.25 |
| | | 30 | 2 | 1.62 ± 0.081 |
| | | 30 | 24 | 1.71 ± 0.13 |
| | | 30 | 49 | 1.64 ± 0.034 |
| | | 30 | 70 | 1.59 ± 0.061 |
| | | 120 | 2 | 1.97 ± 0.46 |
| | | 120 | 24 | 1.84 ± 0.28 |
| | | 120 | 49 | 1.77 ± 0.061 |
| | | 120 | 70 | 1.69 ± 0.062 |

### Supplementary Figures

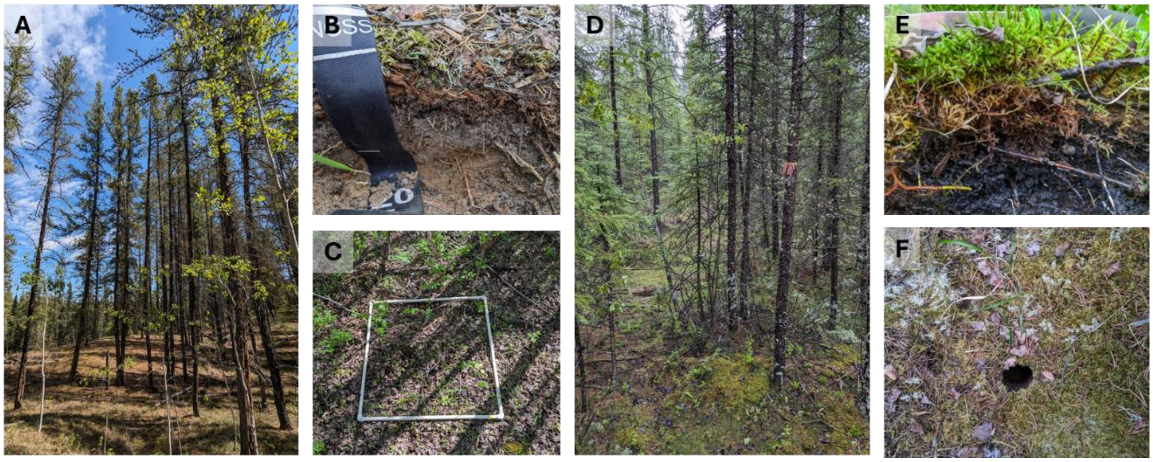

*Figure S1. Photos from Histosol sites showing dominance of spruce trees (A), typical composition of surface layers of soil (B), and an overhead view of the soil surface following core collection (C). Photos from Gleysol sites showing dominance of pine trees (D), typical composition of surface layers of soil (E), and an overhead view of the soil surface (F).*

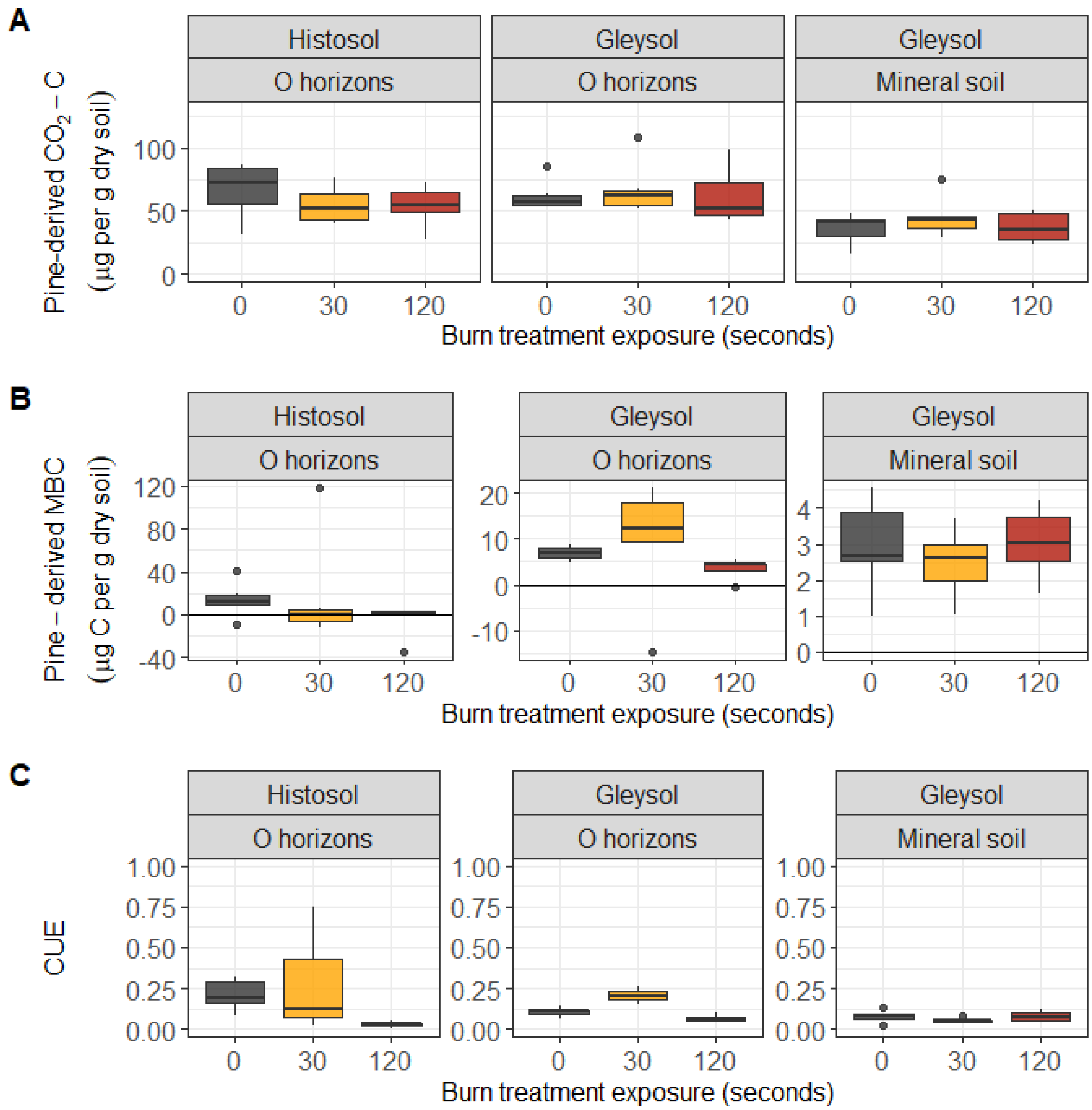


*Figure S2. For completeness, we report results for a parallel pine-specific CUE experiment including (A) pine-derived $CO_2$-C, (B) pine-derived MBC, and (C) pine-specific CUE.*

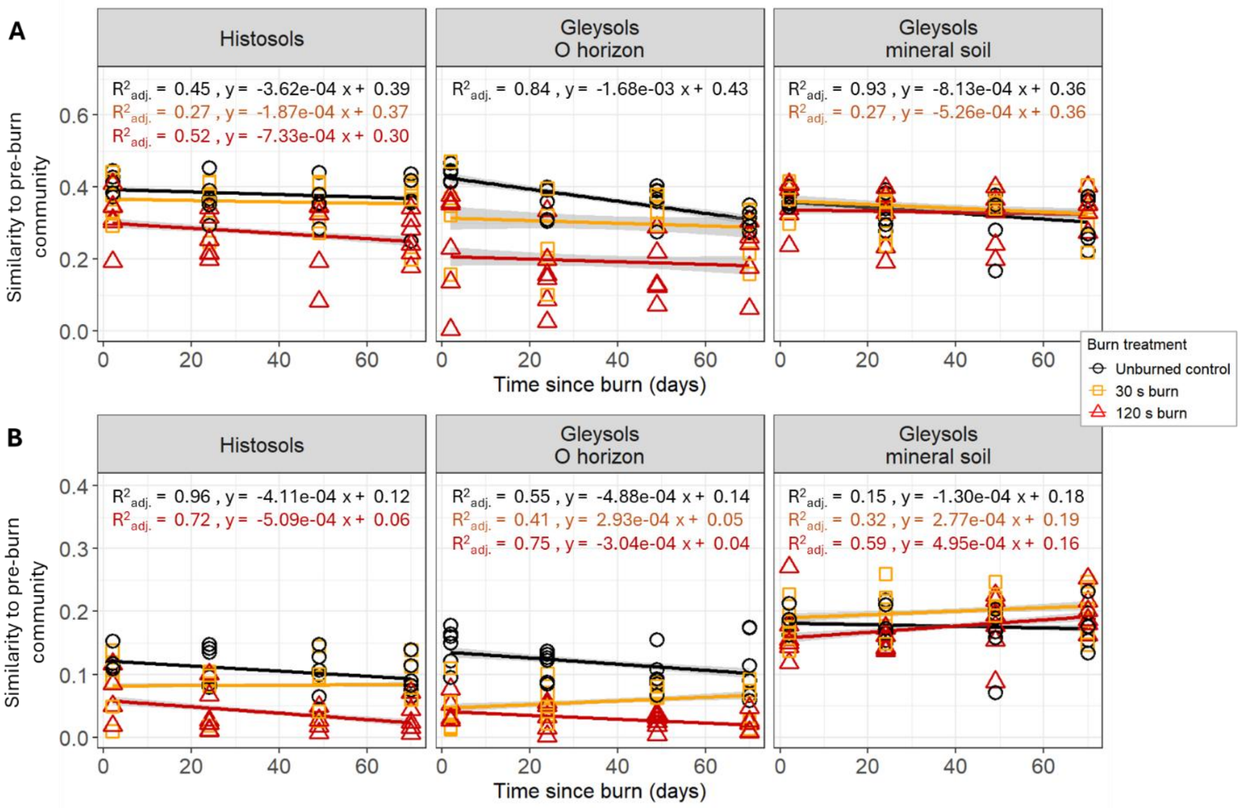


*Figure S3. Average Bray-Curtis similarity of soil (A) bacterial and archaeal communities and (B) fungal communities between a given soil core and all unburned cores within a given soil type and horizon at 2 days post-burn (excluding the control core from the same site). Black, orange, and red represent soils exposed to 0, 30, and 120 s burn treatments, respectively. Linear regressions with 95% confidence intervals (grey shading) are shown to illustrate differences with burn treatment. In instances where the linear regression slope does not differ from zero, best fits are shown, and the slope is not reported.*

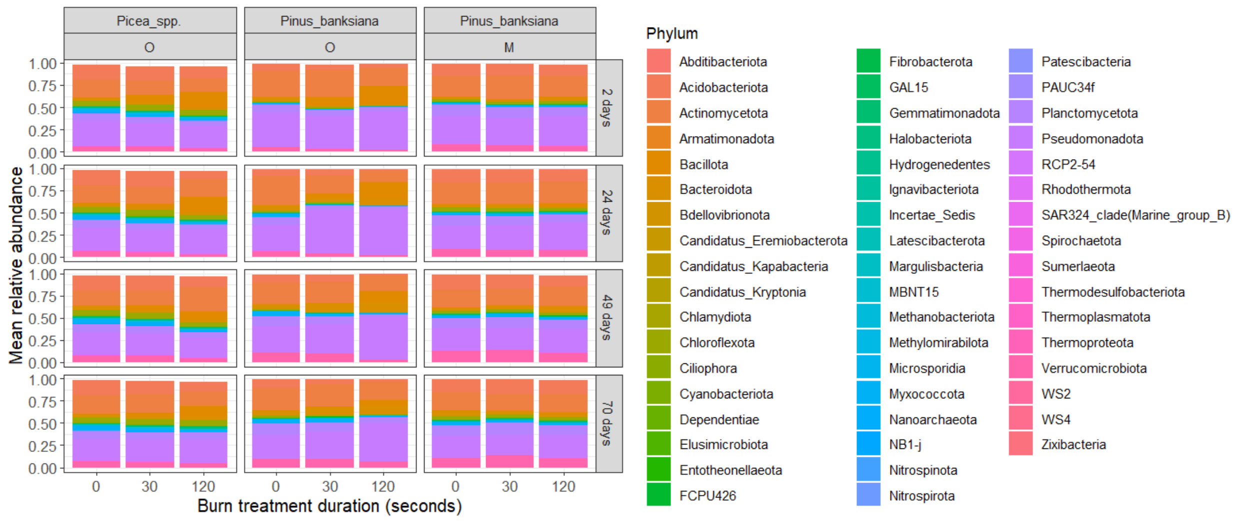


*Figure S4. Mean relative abundance of all bacterial phyla by site type (vegetation and soil horizon), incubation duration (days), and burn treatment duration (seconds).*

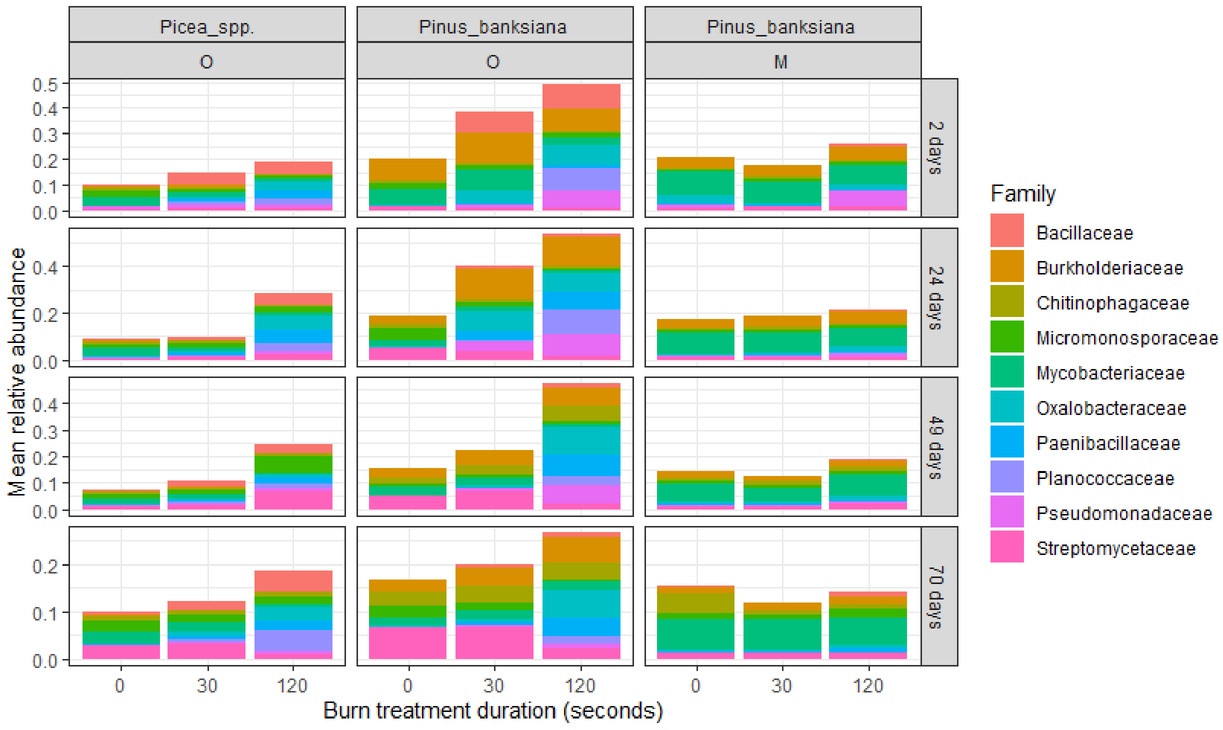


*Figure S5. Mean relative abundance of the 10 most abundant bacterial families by site type (vegetation and soil horizon), incubation duration (days), and burn treatment duration (seconds).*

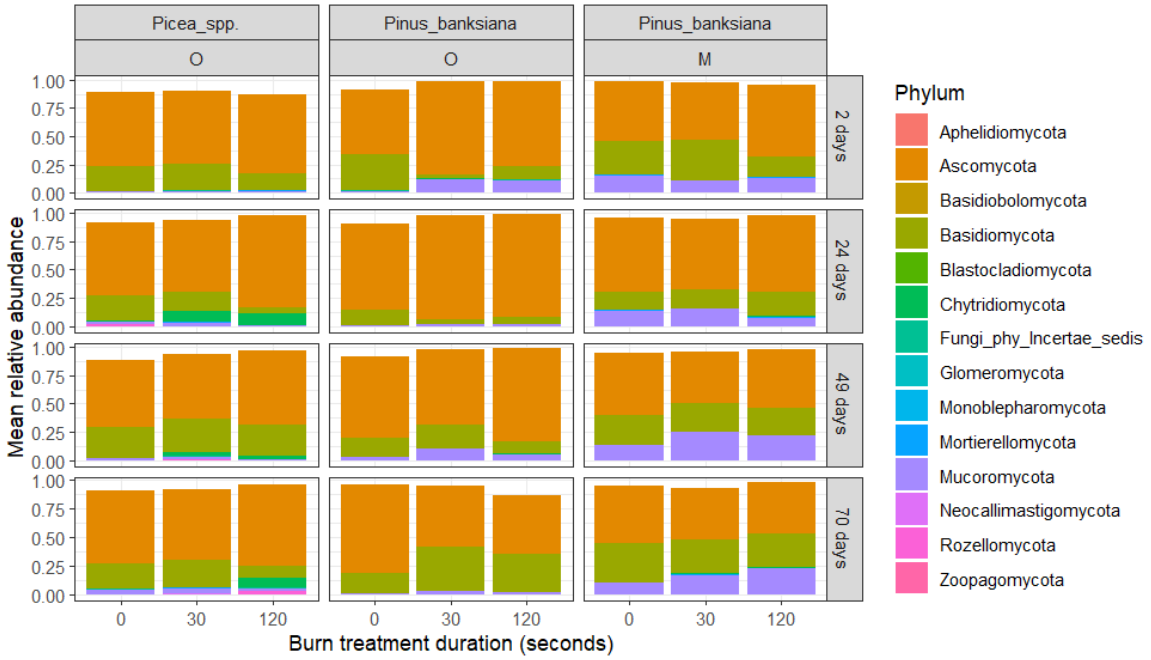


*Figure S6. Mean relative abundance of all fungal phyla by site type (vegetation and soil horizon), incubation duration (days), and burn treatment duration (seconds).*

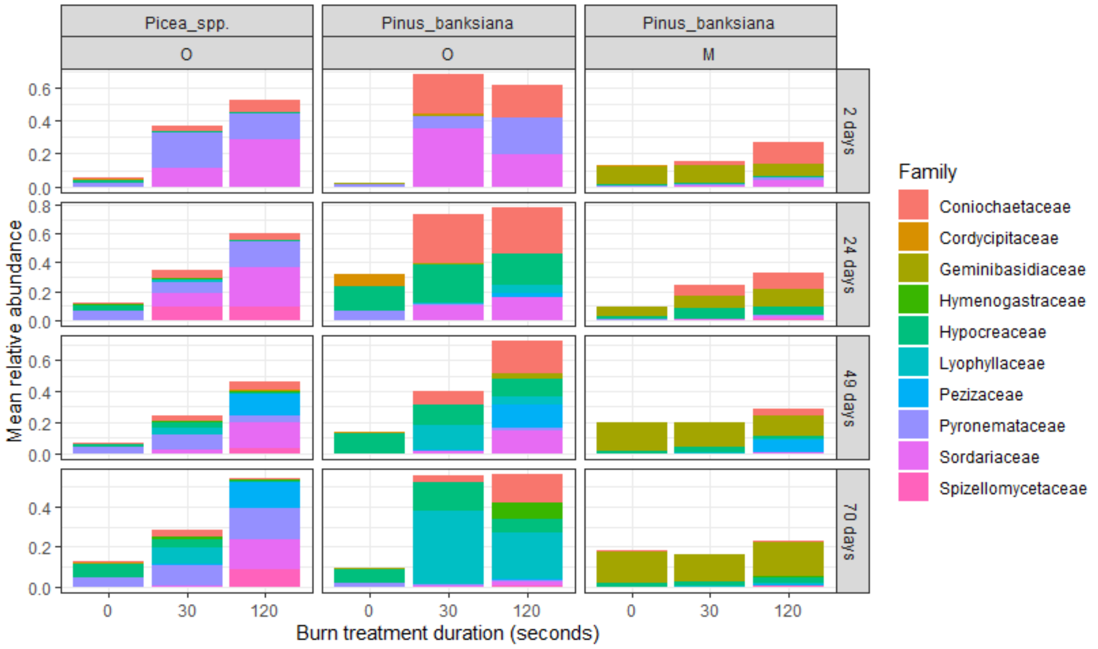


*Figure S7. Mean relative abundance of the 10 most abundant fungal families by site type (vegetation and soil horizon), incubation duration (days), and burn treatment duration (seconds).*

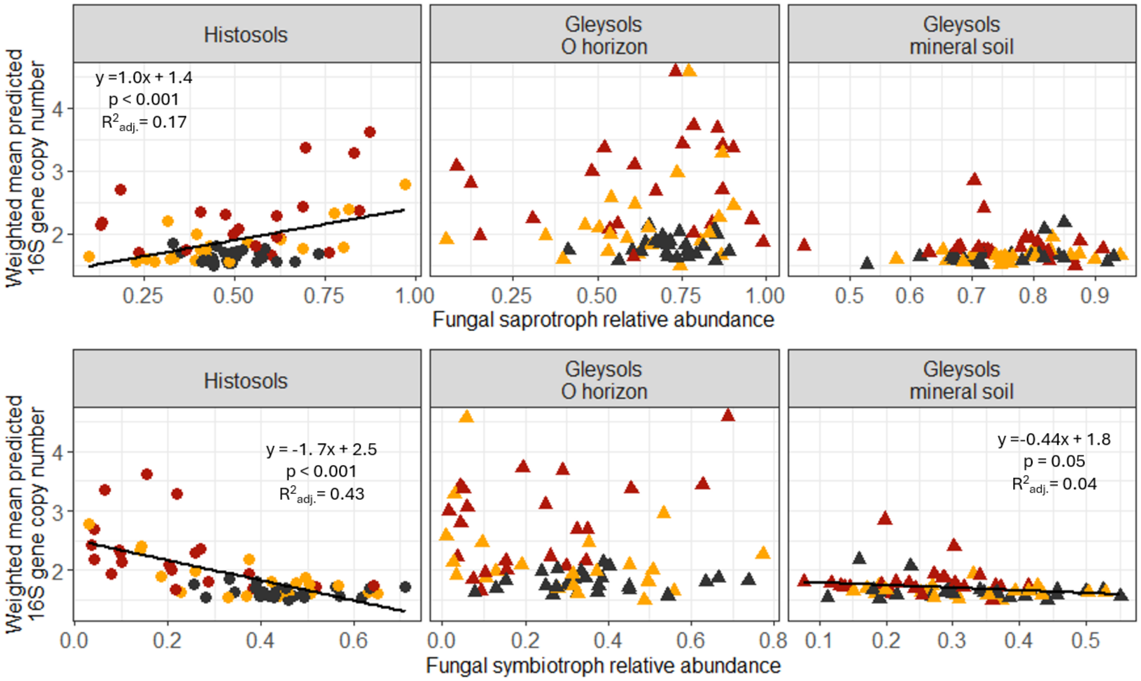


*Figure S8. The relationship between the relative abundance of putative fungal saprotrophs (top panels) and putative fungal symbiotrophs (bottom panels) versus weighted mean predicted 16S rRNA gene copy number in Histosols (left), Gleysol O horizons (middle), and Gleysol mineral soil (right) across all incubation timepoints. Black, orange, and red represent soils exposed to 0 s (unburned controls), 30 s, and 120 s burn treatment durations, respectively. Circles and triangles denote samples from Histosols and Gleysols, respectively.*

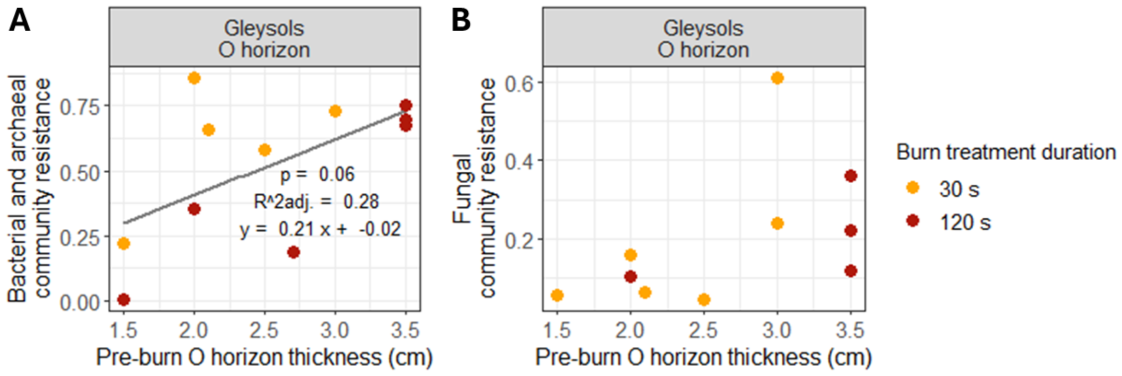


*Figure S9. The relationship between the pre-burn thickness of O horizons in Gleysols and the resistance of bacterial and archaeal communities (left panel) and fungal communities (right panel). Orange and red circles represent soils exposed to the 30 s and 120 s burn treatments, respectively.*

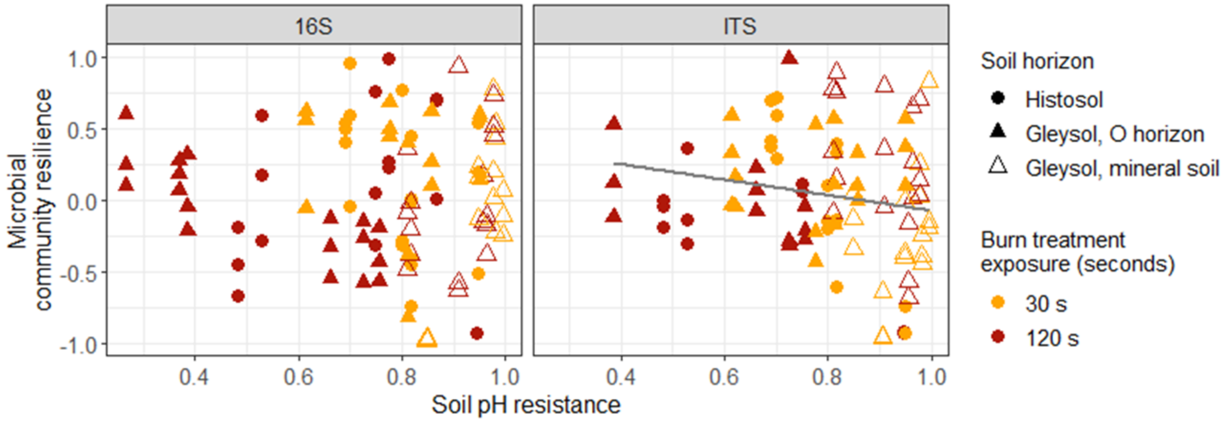


*Figure S10. Relationship between soil pH resistance to burning and bacterial (left panel) and fungal (right panel) community resilience to burning. Orange and red represent soils exposed to the 30 s and 120 s burn treatments, respectively. Circles and triangles represent Histosols and Gleysols, respectively, with filled and unfilled symbols representing organic horizons and mineral soils, respectively.*

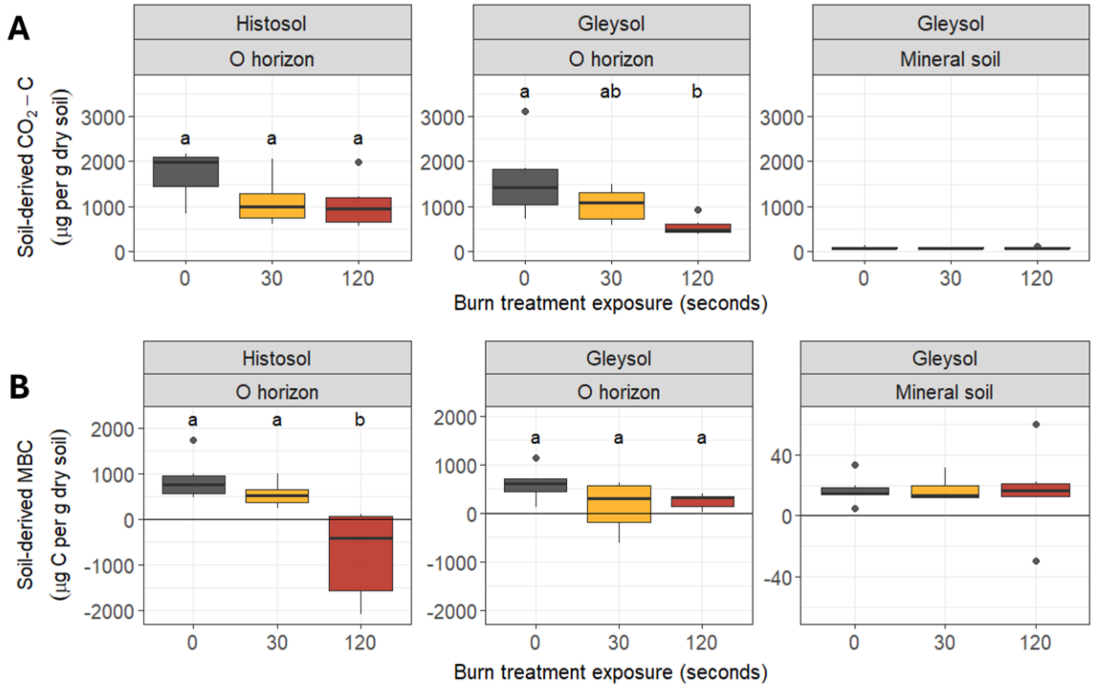


*Figure S11. Soil-derived (A) $CO_2$-C and (B) microbial biomass C in Histosols (left), Gleysol O horizons (middle), and Gleysol mineral soil (right). Black, orange, and red represent soils exposed to 0 s (unburned controls), 30 s, and 120 s burn treatment durations, respectively.*

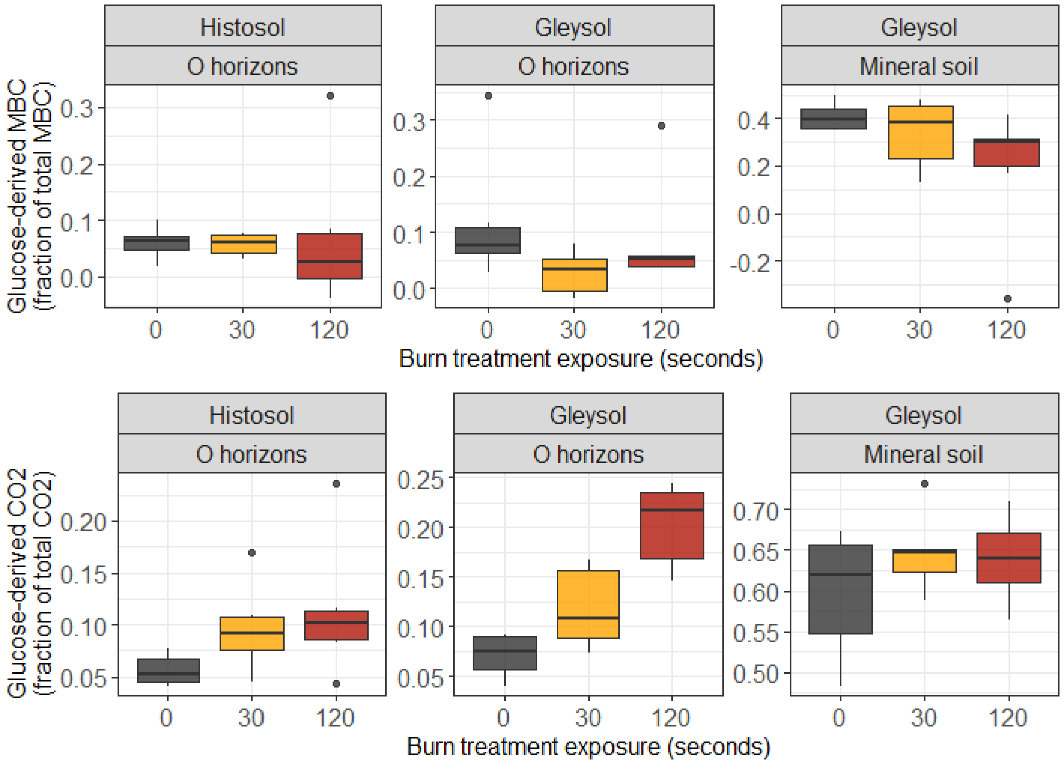


*Figure S12. The effect of burn treatment duration on the fraction of (A) total microbial biomass C and (B) total $CO_2$-C that is glucose-derived in Histosols (left), Gleysol O horizons (middle), and Gleysol mineral soil (right). Black, orange, and red represent soils exposed to 0 s (unburned controls), 30 s, and 120 s burn treatment durations, respectively.*
